# Cellular Mechanisms of Transcranial Magnetic Stimulation in Climbing Fibers and Purkinje Neurons in the Cerebellum

**DOI:** 10.64898/2026.05.12.724125

**Authors:** Yoshio Okada, Chunling Dong, Sergei Makaroff, Padmavathi Sundaram

**Affiliations:** Athinoula. A. Martinos Center for Biomedical Imaging, Department of Radiology, Massachusetts General Hospital, Boston, MA; Division of Newborn Medicine, Department of Pediatrics, Boston Children’s Hospital, Boston, MA 02115; Harvard Medical School, Boston, MA; Worcester Polytechnic Institute, Worcester, MA

**Keywords:** Transcranial Magnetic Stimulation (TMS), Brain stimulation, Neuromodulation, Cerebellum, Activation electric field threshold, Purkinje neurons, climbing fibers

## Abstract

Although transcranial magnetic stimulation (TMS) is widely used for brain stimulation, fundamental issues about its underlying mechanisms remain unresolved. We investigated some of these issues experimentally using an intact isolated turtle cerebellum *in vitro,* employing a novel chamber designed to deliver precisely calibrated induced electric fields along cortical depth. Our results show that single-pulse TMS can directly activate Purkinje cells and climbing fibers, and synaptically activate Purkinje cells via climbing fibers – all within the first 1.2 ms. Specifically, current source density analysis showed that TMS directly (non-synaptically) activated (1) climbing fibers near the bend with intracellular current directed toward the axonal terminals and (2) Purkinje cells directly near the axon initial segment with intracellular current directed toward the distal dendrites. The thresholds for direct activation of climbing fibers and Purkinje cells were found to be very similar, 25 ± 1 V/m. The climbing fibers synaptically activated Purkinje cells, as expected, with intracellular current originating in the proximal dendritic trunk and directed toward the distal dendrites. At higher electric fields (> 58 ± 17 V/m), TMS synaptically activated dendritic currents in Purkinje cells. These results provide new insight into how TMS may activate afferent fibers and cell bodies of cortical neurons.

## INTRODUCTION

Transcranial Magnetic Stimulation (TMS) is a widely used technique for noninvasive stimulation and modulation of human brain function^1,2^. In TMS, a brief (∼400 µs) time-varying current in a coil generates a time-varying magnetic field which in turn induces an electric field (E-field) in the brain tissue, leading to action potentials when the E-field is suprathreshold^1,3–5^. TMS has attracted considerable interest in both basic neuroscience^6^ and clinical medicine due to its ability to non-invasively activate neuronal populations and induce plasticity^7^. It is also FDA-approved for treatment-resistant depression^14^ and migraine^8^ and shows promise in ongoing clinical trials for diagnosing motor system disorders^9–11^ and treating stroke^12–14^ and schizophrenia^15–18^.

While TMS has achieved broad clinical and scientific adoption, the biophysical mechanisms by which it activates neurons remain poorly understood^19^. A point of ongoing debate is whether TMS preferentially activates axons over cell bodies of neurons in the cortex and if so, whether this activation is initiated at axonal bends or terminals^19–22^. It also remains unclear whether TMS can in some settings directly activate neuronal bodies and, if so, where the activation is initiated^19^. In addition, direct empirical measurements of the E-field activation threshold are still not available for specific neurons. Thresholds for activating motor neurons using TMS in the human primary motor cortex (M1) have been estimated to be ∼33-50 V/m using EEG evoked responses^23,24^ and ∼50-70 V/m using electromyography (EMG)^25–27^. Whether these thresholds reflect direct activation of corticospinal motor neurons in M1 or recruitment through synaptically coupled neurons remains unresolved. Simulation studies using realistic neuron models report much higher thresholds (175-430 V/m) for cortical pyramidal neurons^20,22,28–30^. The reasons for this large discrepancy are still unresolved^31^.

These outstanding issues are being actively investigated with *in vivo* and *in vitro* preparations, yet they are still unresolved. Although several *in vivo* animal studies have investigated TMS-induced neuronal activation^32–39^, including some using single-unit recordings^32,36,38,39^, direct estimates of the E-field threshold for activating specific neurons *in vivo* are still lacking. While many *in vitro* studies have been performed^40–55^, few have recorded electrophysiology concurrently with TMS application^40,47,48^. Most *in vitro* TMS studies recorded electrophysiological responses before and after stimulation, limiting observations to post-TMS changes in excitability^44–46,51^. The scarcity of simultaneous TMS and electrophysiology recordings may partly stem from the prolonged TMS artifact that obscures evoked neuronal activity in the crucial period immediately following each TMS pulse^56,57^. In conventional *in vitro* recording chambers, the induced E-field is attenuated within the brain tissue due to the geometry of the experimental setup^56^. Consequently, high intensities on the stimulator are required to activate neurons. But these high-intensity pulses generate prolonged electromagnetic artifacts and mechanical vibrations of the recording electrode, both of which can obscure neuronal activity during the critical period immediately following the pulse.

We used a novel *in vitro* approach that removes some of these major difficulties to address the outstanding issues described above. We used an isolated intact turtle cerebellum since it is well-suited for this investigation due to its orthogonal cellular organization with principal neurons oriented perpendicular to the cortical surface. This geometry enables us to selectively activate a well-defined set of neuronal elements, simplifying the experimental analysis. We developed a chamber design that concentrates the current density and E-field in the tissue, providing strong calibrated E-fields in the tissue at low stimulator intensities, allowing activation of specific neuronal elements with short (< 1 ms) TMS artifacts. We measured the laminar potential profiles with local field potential (LFP) recordings and computed the current-source density (CSD) along the cortical depth in order to infer the origin and direction of the intracellular currents evoked by TMS in the afferent fibers and principal neurons. The novel *in vitro* chamber and the precisely calibrated tissue E-field together allowed us to determine the activation E-field thresholds for both Purkinje cells and climbing fibers.

Our approach uncovered many previously uncharacterized aspects of TMS-induced neuronal activation. Using CSD analysis, we identified how TMS generates intracellular currents in the climbing fibers and Purkinje neurons. Using the calibrated E-fields, we determined that both Purkinje cells and climbing fibers can be activated at very comparable E-field thresholds of ∼25 V/m, slightly lower than those estimated in humans and much lower than those based on simulation studies. As discussed below, our results provide specific new insights into the outstanding issues pointed out above. Our findings may advance understanding of the cellular mechanisms of TMS activation of afferent fibers and cortical neurons.

## RESULTS

### TMS Activates Cerebellar Neurons in a Chamber with a Current Concentrator

In a conventional *in vitro* chamber, the TMS-induced E-field within the tissue is attenuated (Figure S2) by the spread of induced currents throughout the bath and by the effect of the air-saline conductivity boundaries^56^. Achieving a sufficiently strong E-field in the tissue necessitates use of a high stimulus intensity. This, however, produces prolonged artifacts in the electrical recording that obscure signals in the period immediately following stimulation. In order to provide suprathreshold E-fields with short artifacts, we designed a new chamber with a current concentrator to focus the induced current within a cylindrical tunnel containing the cerebellum (Figure 1A). This design is based on earlier studies of effects of applied E-fields on neuronal activation in the cerebellum^58–60^. The TMS artifacts were reduced to ∼1 ms, enabling inspection of evoked neuronal activity immediately after TMS application.

**Figure 1.**
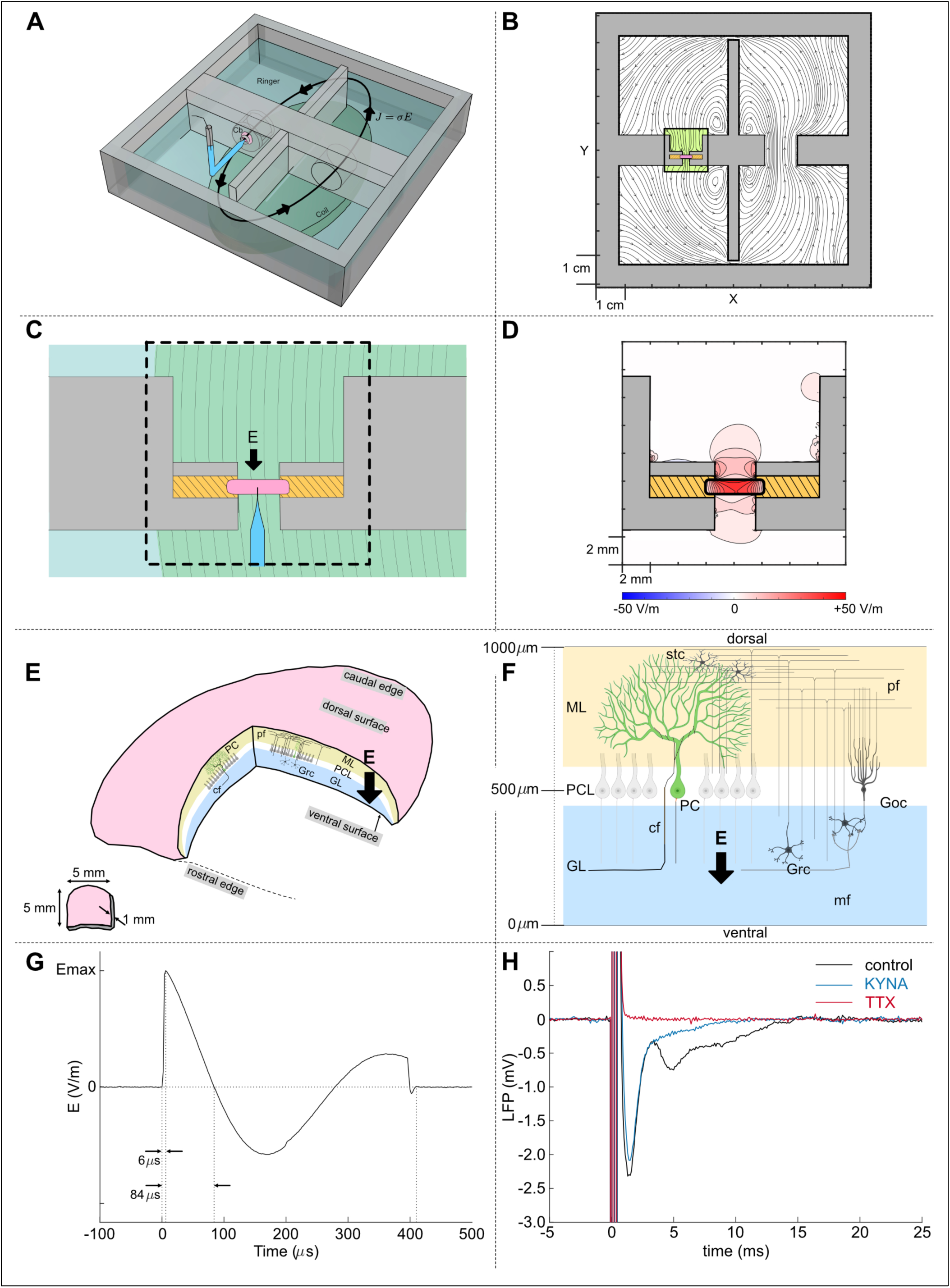
Current concentrator chamber enables strong tissue E-fields at low MSO, facilitating recording of concurrent TMS-evoked LFP in an intact cerebellum *in vitro*. **(A)** Schematic illustration of the recording chamber filled with physiological saline with the intact cerebellum placed in a tunnel with a 3-mm opening. A bent glass micropipette electrode was advanced through the tissue to record the evoked responses. The oval TMS coil was placed below the chamber. When the coil current is oriented in the clockwise direction, the induced current in the bath is oriented in the anti-clockwise direction per Faraday’s law. **(B)** Vector plot of the induced E-field in the saline and in the tissue obtained by computational simulation using BEM-FMM. The chamber design concentrates the current in the tunnel containing the tissue. **(C)** Closeup of tissue placement in the tunnel with inset showing how the E-field was oriented along the Purkinje cells (PCs), pointed from the dendrites towards the soma. **(D)** BEM-FMM simulation of the TMS-induced E-field in the cerebellum showing how the current concentrator design yields strong E-fields at low MSO. At 5% MSO (1.5 A/µs), we expect 41.7 V/m in the center of the cerebellum. **(E, F)** Schematic depiction of the arrangement of the different cerebellar neuronal elements with respect to the applied E-field (ML: molecular layer, PCL: Purkinje cell layer, GL: Granular layer, PC: Purkinje cells, stc: stellate cells, pf: parallel fibers, cf: climbing fibers, Grc: Granule cells, mf: mossy fibers, Goc: Golgi cells). The pf run mediolaterally, and PC, GrC, and Goc run ventrodorsally. **(G)** Waveform of the induced E-field showing the rise time (84 µs) and pulse width (410 µs) measured in saline using a dipole probe. **(H)** Extracellular LFP evoked response to TMS measured 500 µm from the ventral surface in the PC layer in the control condition with normal physiological saline (black), with excitatory synaptic transmission blocked with 10 mM KYNA (blue), and with voltage-gated Na^+^ channels blocked using 2 µM TTX (red). TMS E-field was suprathreshold.

The full experimental setup is shown in Figure S1. An ellipsoidal TMS coil was placed 3 mm below the bottom of the chamber. A biphasic current pulse was applied to the coil using a clinical TMS stimulator (MagPro R30, Magventure, Farum, Denmark). The current pulse generates a time-varying magnetic field which induces an opposing E-field and an accompanying current in the bath according to Faraday’s law^4,61^. Figure 1B indicates the pattern of current flow in the saline bath. Figure 1C provides an expanded view of the current concentrator, with the cerebellum (pink ellipsoid) positioned within the tunnel and surrounded by vacuum grease (yellow). TMS-induced current enters the tunnel from above and exits through the 3-mm-wide tunnel. Boundary element model (BEM) simulation^62,63^ of the conductor geometry shows that current concentration in the tunnel helped induce strong E-fields inside the tissue at relatively low maximum stimulator output (% MSO): 41.7 V/m at 5% MSO (Figure 1D), rather than 13 V/m without the concentrator (Figure S2).

Figure 1E schematically illustrates an intact turtle cerebellum. It is essentially shaped like an ellipsoidal disk of 4-5 mm in diameter. Despite being 1 mm thick, the preparation remains physiologically viable^64,65^. The cerebellar cortex has three layers, consisting of the molecular layer in the dorsal half where the Purkinje cell dendrites are located, the granular layer in the ventral half where the granule cells are located, and the Purkinje cell layer in the middle (500 ± 20 µm).

Figure 1F illustrates the cellular arrangement to aid in interpretation of our results. The cerebellum receives two types of afferent input: the climbing fibers (cf) and the mossy fibers (mf). Climbing fibers enter the cerebellum through the peduncles, course horizontally through the granular layer, then bend to ascend toward the dorsal surface, forming synaptic contacts onto the proximal third of the Purkinje cell dendritic trunk^66–68^. Mossy fibers also enter through the peduncles, course horizontally in granular layer and form synaptic contacts with the glomeruli of the granule cells (Grc). The ascending axons of granule cells bifurcate to produce parallel fibers (pf), which extend approximately 1.5 mm along the mediolateral axis in each direction^66^. The parallel fibers synaptically innervate Golgi cells (Goc). Golgi cells inhibit granule cells at the glomeruli. Parallel fibers make extensive excitatory synaptic contacts with Purkinje cell dendrites in the upper two-thirds of the molecular layer^66,68–70^. Purkinje cells serve as the sole efferent output of the cerebellum, projecting their axons to external targets. These neuronal elements are arranged along three mutually orthogonally axes. In the present study, the induced E-field in the tissue was oriented perpendicular to the cortical surface, as illustrated in Figures 1E and 1F, in order to selectively stimulate the neuronal elements aligned along the depth of the cortex.

Figure 1G shows the time course of the E-field induced by a single biphasic TMS pulse. The E-field reached its peak value 6 µs after pulse application with a rise time of 84 µs and a pulse width of 410 µs. This time course was measured using a dipole probe in the saline bath outside of the current concentrator tunnel^40^. The E-field in the tissue was empirically determined using a calibration procedure described later. This allowed conversion of the strength of the applied stimulus in %MSO to E-field (V/m) in the tissue.

Our setup enabled synchronous activation of large neuronal populations at low stimulator intensities, with responses observed at settings as low as 3-5% MSO. Figure 1H shows an example of the LFP recorded in the Purkinje cell layer produced by a single biphasic TMS pulse with suprathreshold stimulus intensity. In normal physiological saline (black line, Figure 1H), the LFP consisted of an initial spike peaking at 1.2-2.0 ms, followed by a slower component peaking at 4-6 ms. A third component with latency >10 ms was occasionally observed (Figure S3). Kynurenic acid (KYNA) is a non-specific blocker of synaptic transmission mediated by excitatory amino acids (EAAs) and has been shown to effectively block EAA synaptic transmission in turtle cerebellum^60,71,72^. When KYNA (10 mM in this example) was added to the bath, the slower components were eliminated (blue line in Figures 1H and S3). The initial spike remained after KYNA application but was partially reduced; while this reduction was modest in Figure 1H, it was more pronounced in other preparations (Figure S3). Use of calcium-free Ringer solution with replacement of Ca^2+^ with Mn^2+^ also eliminated the slower (4-6 ms) components (Figure S4). Bath application of a Na^+^-channel blocker (tetrodotoxin TTX, 2 µM) abolished all responses except the TMS artifact (red curve in Figures 1H, S3), confirming their neuronal origin.

### Cellular Generators of TMS-Evoked Currents Revealed by CSD Analysis

To identify the generators of the TMS-evoked response, we measured the laminar profile of the LFP and computed the current source density (CSD) along the cortical depth. The laminar profile was obtained by advancing a sharp-tip glass micropipette (< 5 μm tip) along the depth and recording the LFP at each position (N = 5 animals). The laminar profile recorded in a preparation in normal physiological saline is shown in Figure 2A. A single biphasic TMS pulse was applied at t = 0. The artifact during the first 0.9 ms is not shown for clarity. The first component (labelled *a*) peaked at 1.5 ms and showed peak negativity in the granular layer ∼400 µm from the ventral surface. Its polarity reversed in the molecular layer, 700 µm from the ventral surface (marked *). The slower second component at ∼5 ms (labelled *b*) was negative in the granular and Purkinje cell layers with polarity reversal seen around 750 µm (marked *) from the ventral surface.

**Figure 2.**
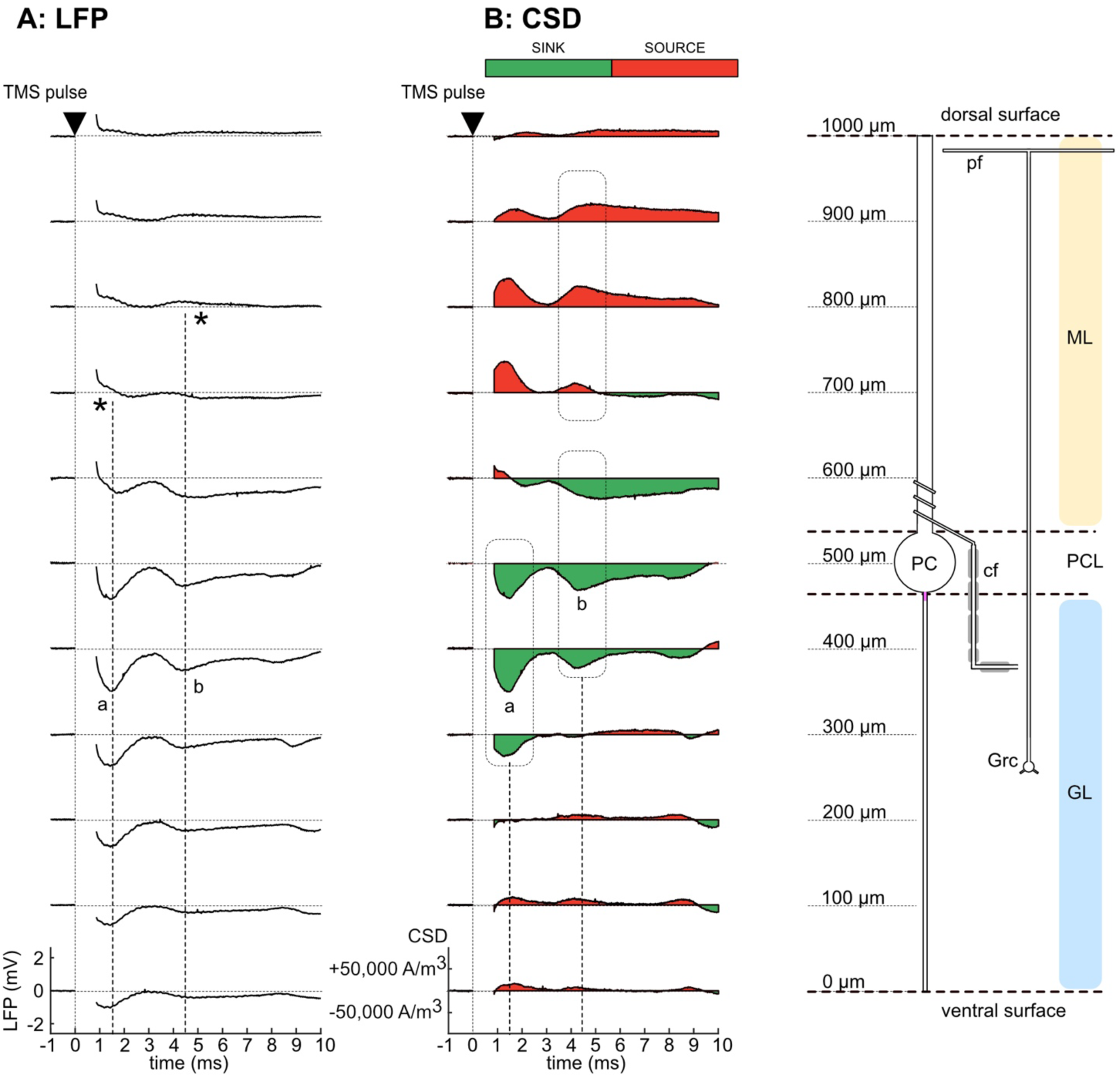
Laminar LFPs and CSD due to a single TMS pulse in normal physiological saline. **(A)** Laminar profile of evoked LFPs (10 ms window post-TMS). TMS stimulation was suprathreshold. The recording depths are indicated in µm as distance from the ventral surface. Note the initial spike “a” (1.2-2.0 ms latency, negative in the GL, peaking 300-500 µm with a polarity reversal at 600-700 µm) and the second slower component “b” (5.0 ms latency, also negative in the GL and a polarity reversal at ∼750 µm). The TMS stimulus artifact ending at 0.9 ms is not shown. **(B)** CSD profile of the responses in (A) assuming tissue conductivity of 0.2 S/m. The initial spike “a” had a current sink (green) in the GL and the PCL and corresponding source (red) in the ML. The CSD profile of the 5 ms component (“b”) was similar but shifted dorsally compared to the spike. Cell layers are indicated on the right (ML: molecular layer, PCL: Purkinje cell layer, GL: Granular layer). PC: Purkinje cell, Grc: Granule cell, cf: climbing fiber, pf: parallel fiber. Data shown are for one preparation.

We used CSD analysis to estimate the underlying transmembrane current density from the second spatial derivative of the laminar profile^73,74^. CSD analysis establishes the spatial distribution of extracellular current sources and sinks in the tissue. Current sinks indicate where current enters neurons from the extracellular space, while current sources indicate where current exits neurons into the extracellular medium^73^. The CSD profile corresponding to the laminar potential data in Figure 2A is shown in Figure 2B. The initial spike had a sink located in the granular and Purkinje cell layers (300-500 µm) with a corresponding source in the molecular layer. This CSD pattern indicates that the underlying intracellular current was directed dorsally for the spike. The slower 5 ms component had a sink that peaked in the Purkinje soma and proximal dendritic trunk regions (500-600 µm from the ventral surface) with a corresponding source in the molecular layer. The underlying intracellular current was directed toward the dorsal surface. Figure 3 shows the laminar potential profile (A) and CSD profile (B) for the same cerebellar preparation as in Figure 2 with synaptic transmission blocked with 10 mM KYNA. The synaptic block reduced but did not eliminate the initial spike but abolished the slower 5 ms component. This indicates that the initial spike was at least in part non-synaptic in origin, while the slower component was completely synaptically mediated. All components were eliminated by 2 µM TTX (Figure S5), showing that they were neuronal in origin.

**Figure 3.**
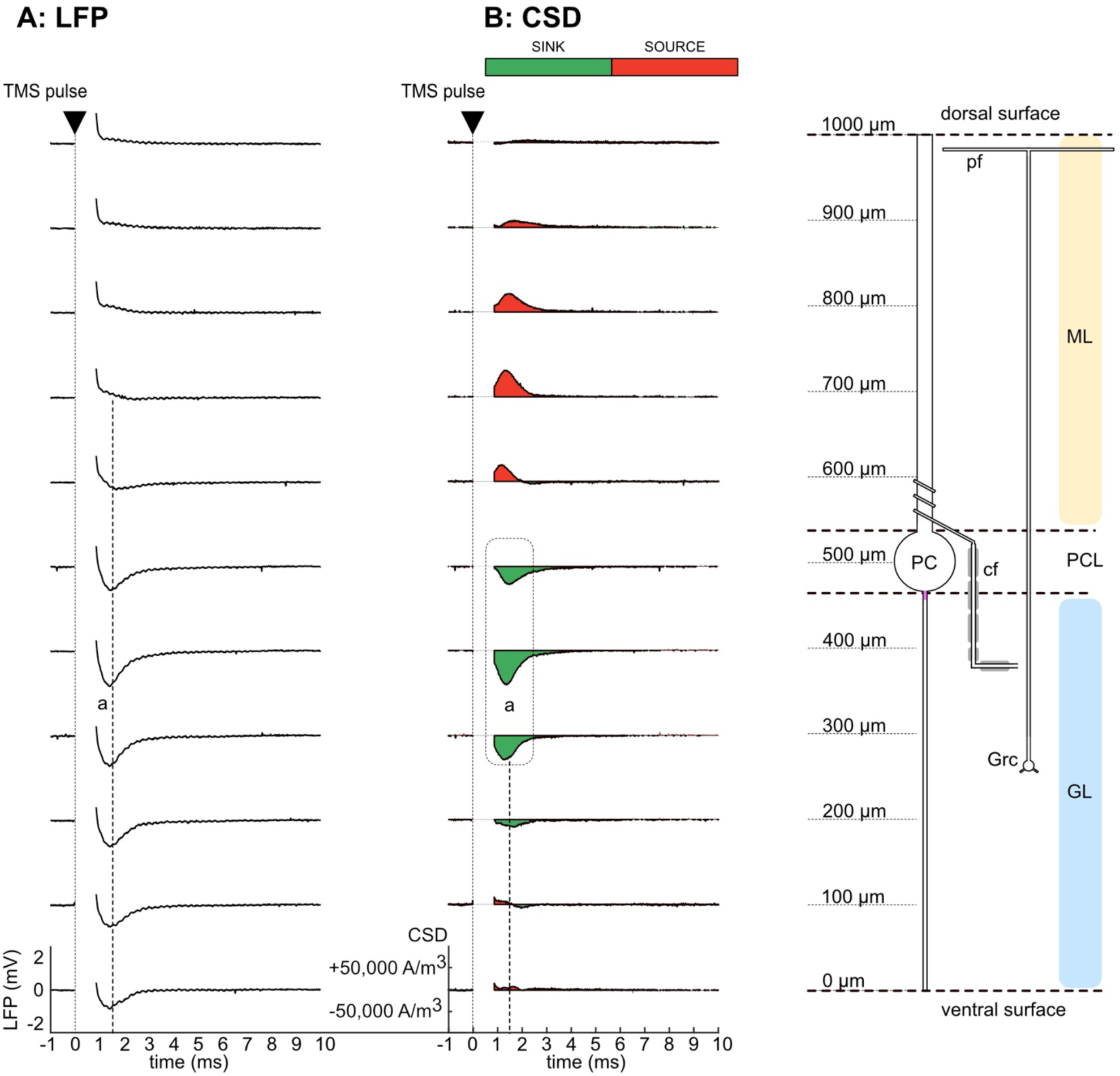
Laminar LFPs and CSD due to a single TMS pulse with excitatory synaptic transmission blocked (10 mM KYNA). **(A)** Laminar profile of evoked LFPs (10 ms window post-TMS). The TMS stimulation was suprathreshold. The recording depths are indicated in µm as distance from the ventral surface. Note that the initial spike “a” (1.2-2.0 ms latency) remains suggesting that it was mostly non-synaptic in origin while the later 5 ms component was blocked by KYNA suggesting that it was synaptic in origin. The TMS stimulus artifact ending at 0.9 ms is not shown. **(B)** CSD profile of the responses in (A) assuming tissue conductivity of 0.2 S/m. The initial spike “a” had a current sink (green) in the GL and the PCL and corresponding source (red) in the ML. Cell layers are indicated on the right (ML: molecular layer, PCL: Purkinje cell layer, GL: Granular layer). PC: Purkinje cell, Grc: Granule cell, cf: climbing fiber, pf: parallel fiber. Data were measured in the same preparation as in Fig. 2.

To identify the neural generators of the CSD sink-source patterns, we examined the CSD profiles as a function of depth in normal (CTRL, Figure 4A) and KYNA saline (KYNA, Figure 4B) averaged across N=5 preparations. The shaded region around each average CSD profile represents the standard error of mean (±1 SEM) across all animals. As indicated in the inset, the initial spike peaked at ∼1.2 ms, but the spike lasted for more than 2 ms. We therefore analyzed the CSD profiles at three latencies: 1.2 ms (blue dot) and 2.0 ms (red dot), representing the initial spike, and 5.0 ms (green dot), near the peak of the slower wave. Figure 4A shows the average CSD profiles for the 1.2 ms and 2.0 ms components in CTRL saline with intact synaptic transmission. The location of current sink peak shifted from the granular layer (400 µm) to the Purkinje cell layer (500 µm) between the 1.2 ms and 2.0 ms components. This shift was significant according to ANOVA (latency x depth interaction significant at p = 0.05, F(10,40) = 14.8). Figure 4B shows the CSD profiles in KYNA with excitatory synaptic transmission blocked. A similar shift in the location of the current sink is observed in Figure 4B between the 1.2 ms and 2.0 ms components and was found to be statistically significant according to ANOVA (latency x depth interaction significant at p = 0.05, F(10,40) = 4.10). These significant shifts in both CTRL and KYNA indicate that the peak of current sink at 1.2 ms was more ventral than the 2.0 ms component, located in the granular layer. Since the current sink peaked in the granular layer with the corresponding source in the molecular layer, we infer that TMS activated the climbing fibers near the bend in the granular layer, producing intracellular currents directed toward the axonal terminals. This direction differs from that predicted by some biophysical modeling studies, which propose that TMS preferentially depolarizes axons at their terminals^19–22^. While the current sink for the 1.2 ms component peaked in the granular layer, the 2.0 ms sink peaked in the Purkinje cell layer, with a corresponding source in the molecular layer. This sink-source pattern suggests that the 2.0 ms component predominantly reflects currents originating in the somatic region of Purkinje cells and directed toward the distal dendrites.

**Figure 4.**
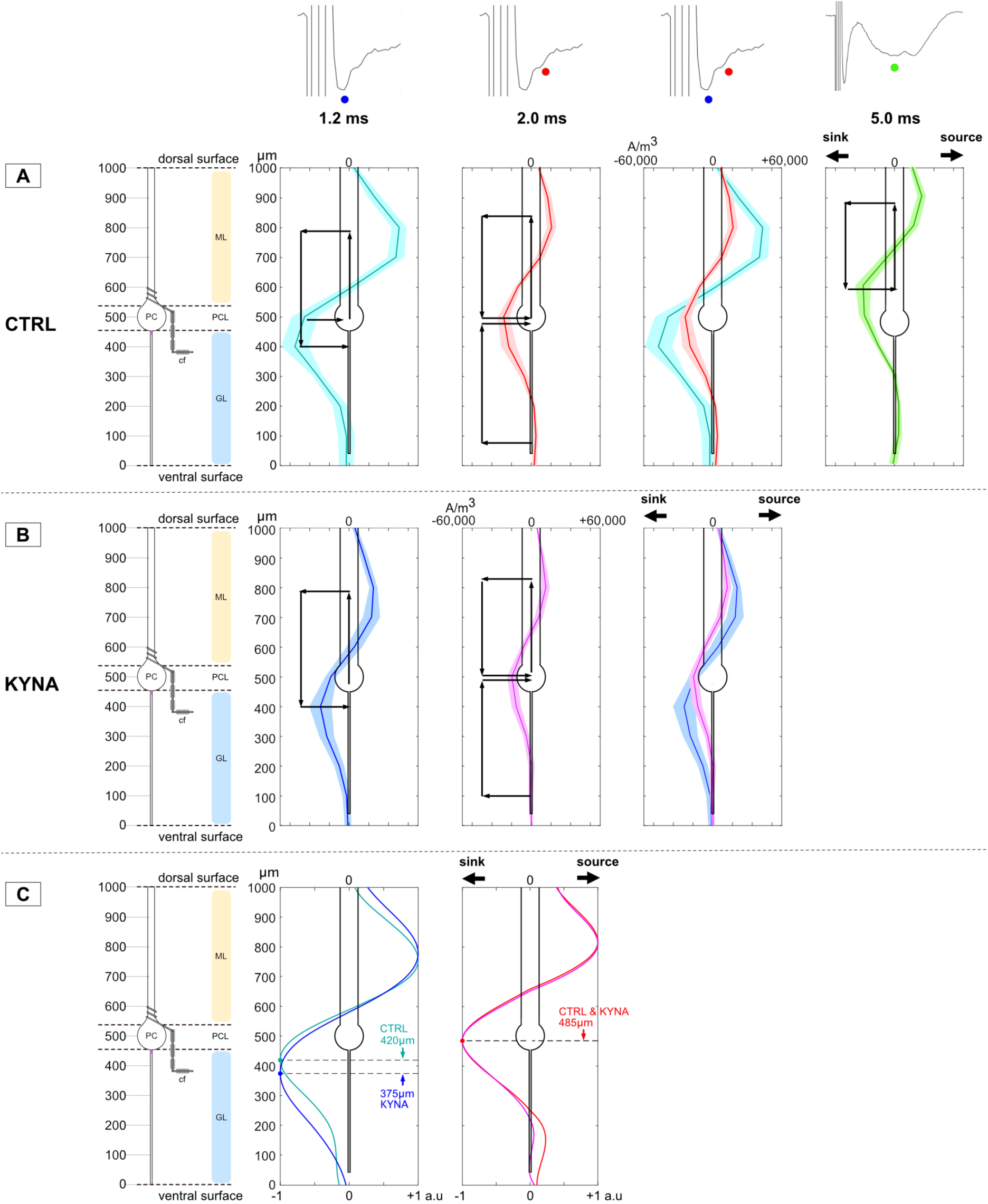
Sink-source patterns in averaged CSD profiles for 1.2 ms, 2.0 ms and 5.0 ms components in normal and KYNA saline indicate underlying neural generators. (A) Average CSD (N=5) profile for different response components due to a single TMS pulse in normal saline with synaptic transmission present. The 1.2 ms component had a peak current sink at 400 µm in the granular layer while the 2.0 ms component had a peak current sink at 500 µm in the Purkinje cell layer. For the 5.0 ms component, the peak current sink was at 600 µm. (B) Average CSD (N=5) profile for different response components due to a single TMS pulse with synaptic transmission blocked (10 mM KYNA). The 1.2 ms component had a peak current sink at 400 µm in the granular layer while the 2.0 ms component had a peak current sink at 500 µm in the PC layer. Both components were affected by KYNA suggesting that they both had synaptic and non-synaptic generators. (C) Interpolated, normalized CSD profiles for 1.2 ms, 2.0 ms components are overlaid to inspect the shift in the current sink due to the KYNA synaptic blockade for the 1.2 ms (CTRL->KYNA: 420 μm->375 μm) and the 2.0 ms component (CTRL->KYNA: no change (485 μm)).

The current sinks and sources corresponding to the 1.2 ms and 2.0 components were both significantly reduced by KYNA application (paired t-test: p < 0.04 (1.2 ms), p < 0.05 (2.0 ms)). The reduction of both components following synaptic blockade indicates that each was partially driven by synaptically mediated currents. Since the spike at 1.2 ms appears to be partially generated by climbing fiber activation, we hypothesize that the synaptic contributions to both 1.2 ms and 2.0 ms spike components are due to climbing fiber synaptic inputs onto the Purkinje cell dendrites. This interpretation is supported by the broad spatial extent of the 2.0 ms current sink, which spanned up to 600 μm (Figure 2B). This spatial extent corresponds to region containing the proximal dendritic trunk of Purkinje cells, where climbing fibers are known to make synaptic contact.

The CSD profile of the slower 5.0 ms component (Figure 4A, right; green dot in the inset) had a peak current sink at 600 μm depth from the ventral surface, in the region of the proximal dendritic trunk. KYNA blocked this component (Figures 1H and 3). These results indicate that the slow wave was synaptically generated by climbing fiber activation of the proximal dendritic trunk of Purkinje cells with the intracellular currents directed toward the distal dendrites.

Figure 4C provides additional details regarding the underlying generators of the spike at 1.2 and 2.0 ms. This figure shows the CSD profiles obtained by interpolating and normalizing the empirical profiles in Figures 4A and 4B. For the 1.2 ms component, Figure 4C (left) shows that KYNA application shifted the peak current sink from 420 μm in CTRL toward the granular layer (375 μm) in KYNA. The two CSD profiles for the 1.2 ms component differed significantly between the CTRL and KYNA conditions; ANOVA showed a significant treatment x depth interaction (F(10,40) = 7.96, p < 0.005, Greenhouse-Geisser corrected). This shift supports the hypothesis that climbing fibers contributed significantly to the 1.2 ms spike. By eliminating synaptic activation of Purkinje cells, the KYNA condition increases the relative contribution of the climbing fibers, leading to a larger shift of the peak toward the granular layer as observed.

The CSD profiles at 2.0 ms were nearly identical in CTRL and KYNA, with the peak current sink at the same location (485 μm). There was no significant treatment x depth interaction (F(10,40) = 3.70, p > 0.05 with Greenhouse-Geisser correction). The virtually identical CSD profiles indicate that the spike at 2.0 ms was generated primarily by Purkinje cells, with minimal climbing fiber contribution. Had the climbing fibers contributed to the CSD profile, the peak current sink would have shifted toward the granular layer in KYNA as for the 1.2 ms component. This conclusion is consistent with the current sink peaking in the Purkinje cell layer (485 μm) in both conditions, as expected if the climbing fiber contribution is negligible. By contrast, the peak of the 1.2 ms component was shifted slightly toward the granular layer (420 μm) in the CTRL condition, indicating some contribution of the climbing fibers.

### Determination of E-field Activation Threshold for the Initial Spike and Slow wave

To determine the activation threshold for climbing fibers and Purkinje cells, we first determined the thresholds for activating the spike and the slow wave, then linked these findings with the CSD analysis to estimate the corresponding E-field thresholds for climbing fibers and Purkinje cells.

As the first step, the E-field induced in the tissue was calibrated by measuring the electric potential in the saline and in the tissue at a fixed intensity (5% MSO, 90 V). This was done by advancing a bent sharp-tip glass micropipette horizontally along the axis of the current concentrator tunnel and measuring the electric potential produced by the TMS pulse (inset at bottom of Figure 5A). The pipette tip was coated with Sylgard to reduce electrode capacitance and increase recording bandwidth. This allowed us to accurately measure the peak electric potential 6 µs after TMS onset. Figure 5A shows the calibration results for 5 preparations. In all cases, the potential increased slowly through the saline and then steeply upon entering the cerebellum via the ventral surface. The E-fields in saline and tissue were estimated from the slope of a least-squares linear fit to the measured potential profiles in each region, at the fixed stimulus intensity. The calibration measurements showed that the E-field at 5% MSO ranged between 30.5 and 40.6 V/m (35.6 ± 4.2 V/m) in the tissue and between 5.3 and 7.5 V/m (6.3 ± 0.87 V/m) in the saline. The ratio of the average E_tissue_ to E_saline_ was 5.6. This agrees with conductivity ratios of saline (1.33 S/m) to tissue (0.15 S/m in the granular layer, 0.25 S/m in the molecular layer) measured in a quasistatic E-field^75^.

**Figure 5.**
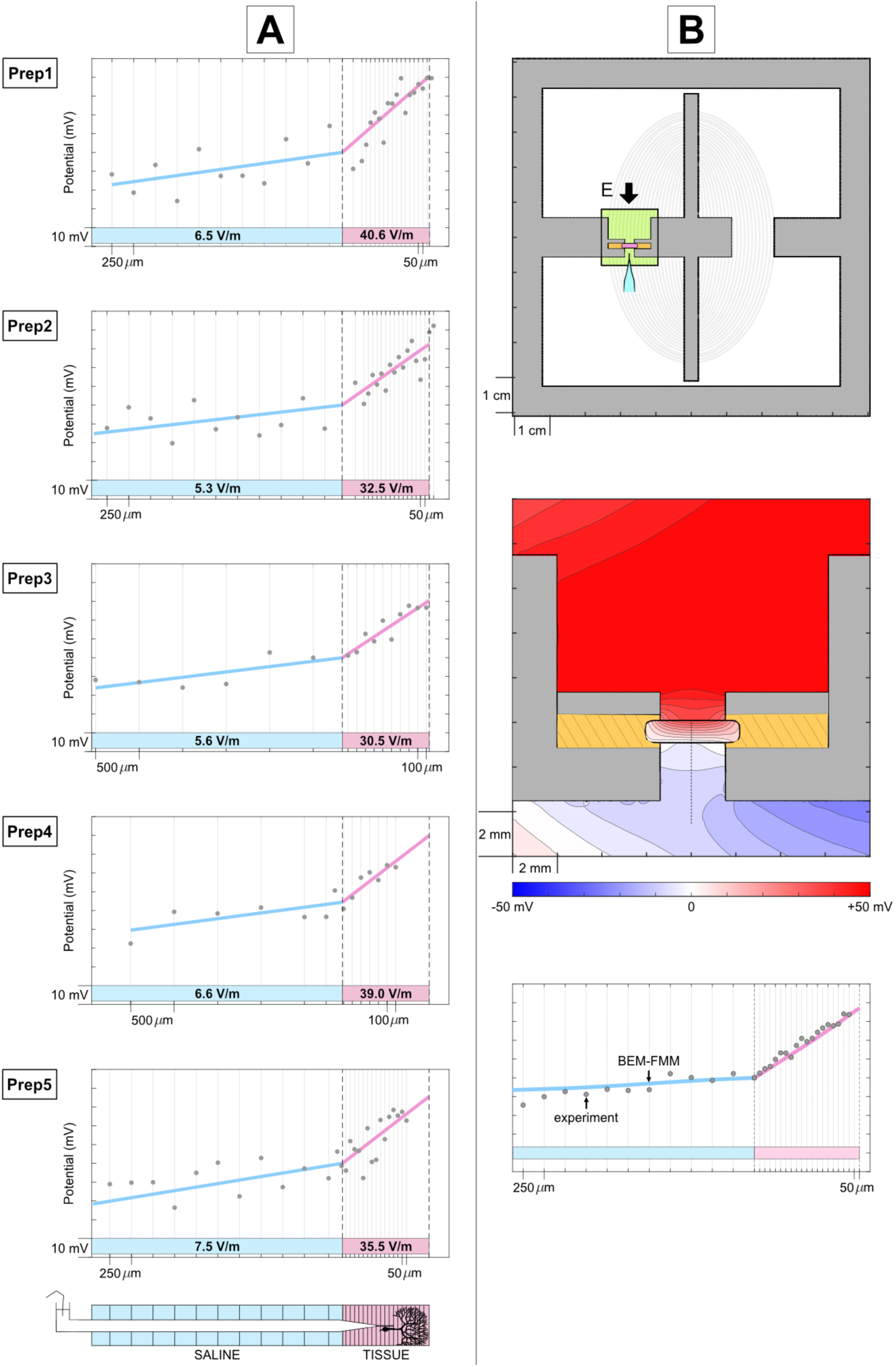
Electric Field Calibration to Determine TMS Activation Thresholds. **A.** To convert the horizontal axis from % MSO to V/m, we calibrated the recording chambers in each of the N=5 cases at fixed MSO (5%). The line profile of the electric potential was measured every 250 µm in the saline (blue) and every 50-100 µm in the tissue (pink). This yielded a calibration factor (MSO(%) -> Tissue E-field) for each preparation. **B.** Top: Realistic model of the tissue in the current concentrator chamber for BEM simulation. Middle: Predicted electric potential in the tissue and saline due to TMS at 5% (from BEM). Bottom: Good agreement (R^2^ = 0.96) between the predicted electric potential profile to the measured electric potential profile in saline and tissue.

We validated our E-field measurements by comparing the measured electric potentials to theoretical values computed using boundary element model (BEM) simulation of the current concentrator chamber (Figure 5B(i))^62,63^. Figure 5B(ii) shows the theoretical electric potential distribution obtained from BEM in the region-of-interest highlighted in the top; as in the experiment, this simulation corresponds to 5% MSO (1.5 A/µs). The empirical and theoretical potential profiles are overlaid in Figure 5B(iii). The average E-field in the cerebellum based on measured potentials was 35.6 V/m, compared to 41.7 V/m obtained using the BEM-FMM. The proportion of the variance in the data accounted for by the BEM-FMM was high (R^2^ = 0.96). The close match provides validation of the empirical estimates.

Following calibration, we determined the E-field thresholds for the 1.2 ms and 2.0 ms spikes, and the 5.0 ms wave. To do this, we recorded the LFP in the Purkinje cell layer (at 500 µm depth) as a function of stimulus intensity (2-15% MSO). Figure 6A shows an example of the LFP measured in one preparation (same preparation as Figure 5A, Prep 1) with synaptic transmission intact and blocked. The stimulus intensities were converted to E-fields in the tissue using the calibration factor specifically for that preparation (5% MSO => 40.6V/m, from Figure 5A, Prep 1).

**Figure 6.**
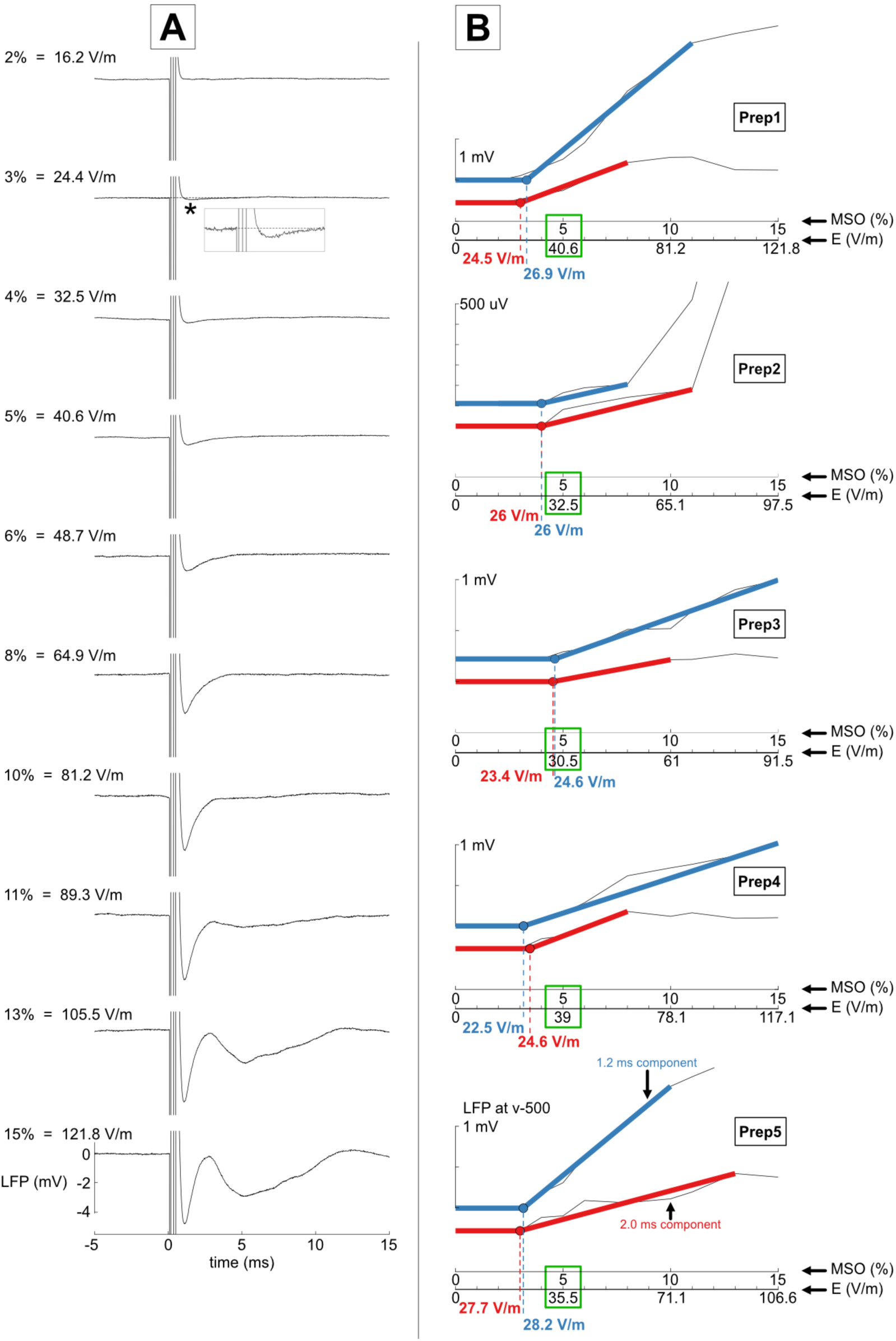
Determination of E-field Threshold for the Initial Components in N=5 preparations. **A.** LFP recorded in the Purkinje cell layer (500 µm depth) as a function of stimulus intensity (2% to 15% MSO). **B.** For N=5 preparations, plots show the LFP amplitude for the 1.2 ms (blue) and 2.0 ms (red) components as a function of TMS stimulus intensity (MSO %). The top plot (Prep1) corresponds to the same preparation as Figure 6A, with the calibration factor of 5% -> 40.6 V/m. The calibration factors show expected experiment-to-experiment variability and allow determination of the thresholds in each case. The average thresholds for the 1.2 ms (25.6 ± 2.2 V/m) and 2.0 ms (25.3 ± 1.6 V/m) components were nearly identical.

Figure 6B shows the amplitude of the 1.2 ms (blue) and 2.0 ms (red) components as a function of stimulus intensity for each of the 5 preparations. Preparation-specific calibration factors (Figure 5) were used to convert the % MSO to corresponding E-field in the tissue. Least-squares lines were fit to the subthreshold and suprathreshold LFP data for each component, with their intersection defining the activation threshold. These thresholds (Figure 6B) were: 22.5-28.2 V/m for the 1.2 ms component and 23.4-27.7 V/m for 2.0 ms component.

Figure 7 shows the E-field thresholds for all three components (1.2 ms, blue; 2.0 ms, red; 5.0 ms, green) averaged across the same 5 preparations. The LFPs were recorded as a function of %MSO and converted to tissue E-field (V/m) using the empirically determined preparation-specific calibration factors (Figure 5). Average LFPs were obtained by plotting the individual data along the common E-field axis. Figure 7 shows the average LFPs at 500 μm depth for all three components with shaded areas representing the standard deviation. The thresholds - 25.6 ± 2.2 V/m (1.2 ms), 25.3 ± 1.6 V/m (2.0 ms) and 58.1 ± 17 V/m (5 ms) - are shown as dashed arrows, with the horizontal bar indicating the standard deviation across 5 preparations. The 1.2 ms and 2.0 ms spike thresholds were nearly identical, whereas the threshold for the subsequent wave was much higher.

**Figure 7.**
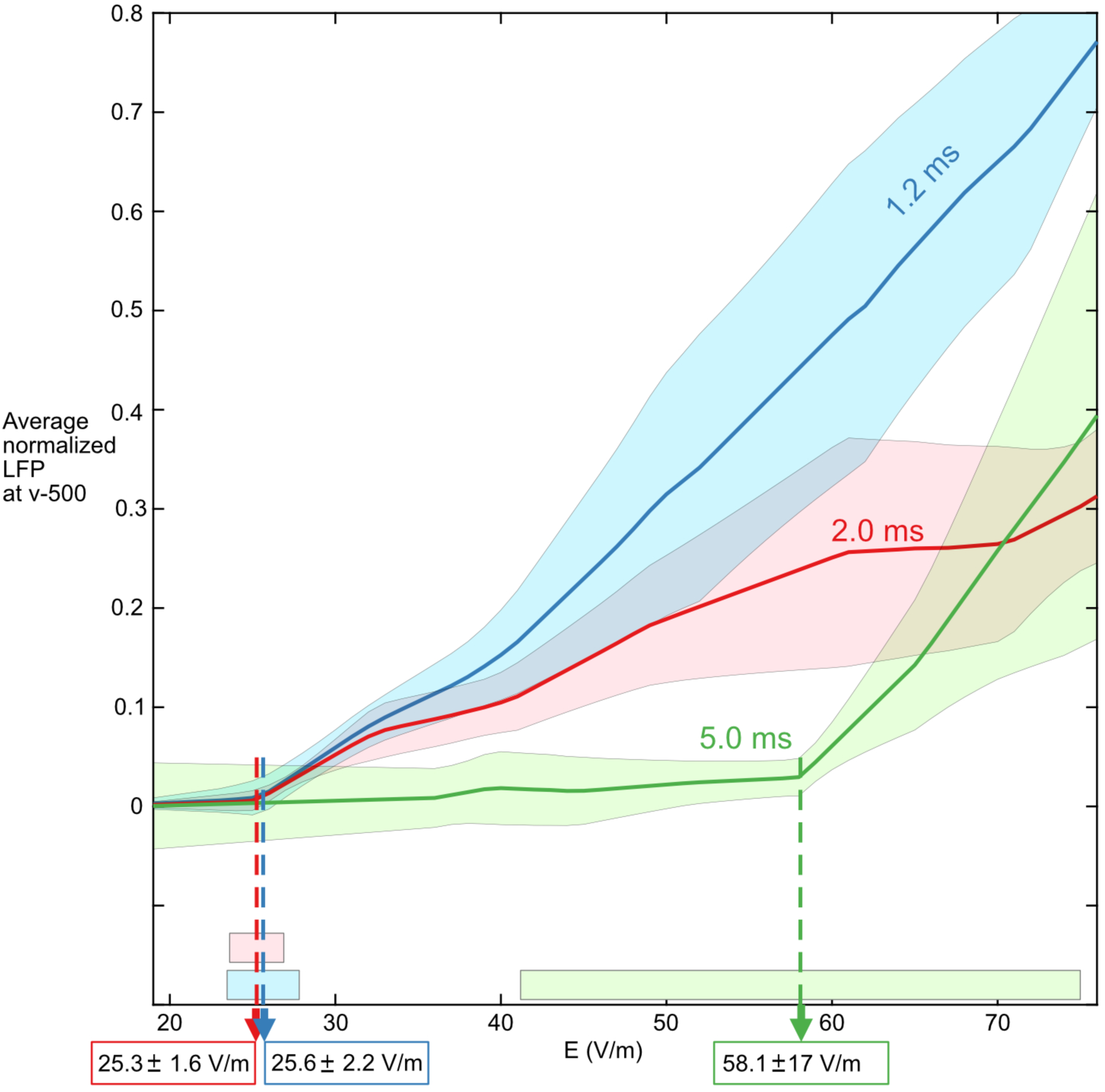
TMS activation thresholds for the 1.2 ms, 2.0 ms and the 5 ms components in normal saline. Average (N=5) peak LFP values for the 1.2 ms, 2.0 ms and 5.0 ms components as a function of measured E-field in the tissue. The average thresholds were 25.6 V/m (1.2 ms), 25.3 V/m (2.0 ms) and 58.1 V/m (5 ms). The thresholds for the 1.2 ms and 2.0 ms components were statistically the same. The 5 ms component had a much higher threshold.

Figure 8A shows an example of the E-field threshold curves for the 1.2 ms and 2.0 ms components in CTRL and KYNA for a single preparation. LFPs were recorded in the Purkinje cell layer (500 µm) across the full %MSO range in both CTRL and KYNA (10-14 trials per stimulus intensity for each preparation; 5 preparations). The %MSO was converted to tissue E-field using the mean threshold value from Figure 7 (25.5 V/m). Spike thresholds for the 1.2 ms and 2.0 ms components were estimated in both CTRL and KYNA conditions by performing least-squares line fits to the suprathreshold LFP amplitudes for each condition. The group average thresholds (Figure 8B) were: CTRL 1.2 ms - 26.2 ± 0.6 V/m; CTRL 2.0ms - 25.5 V/m; KYNA 1.2 ms - 25.5 ± 0.9 V/m; KYNA 2.0 ms - 25.4 ± 1.0 V/m. A two-way repeated measures ANOVA revealed no significant main effect of solution (CTRL vs. KYNA; *F*(1,4) = 0.28, *p* = 0.623), or latency (1.2 ms vs. 2.0 ms; *F*(1,4) = 3.46, *p* = 0.136), and no significant solution × latency interaction (*F*(1,4) = 0.001, *p* = 0.976). Thus, these four thresholds were statistically indistinguishable. The slope was steeper for CTRL than KYNA at 1.2 ms (0.232 ± 0.051 vs. 0.160 ± 0.037, mean ± 1 SEM) and the difference was statistically significant (p <0.02, repeated measures Student’s t-test). The corresponding slopes at 2.0 ms were 0.135 ± 0.019 for CTRL and 0.131 ± 0.017 for KYNA. The difference between the paired slopes at 2.0 ms was not significant (p = 0.05). The significant difference in slope at 1.2 ms indicates a synaptic contribution to the LFP in CTRL.

**Figure 8.**
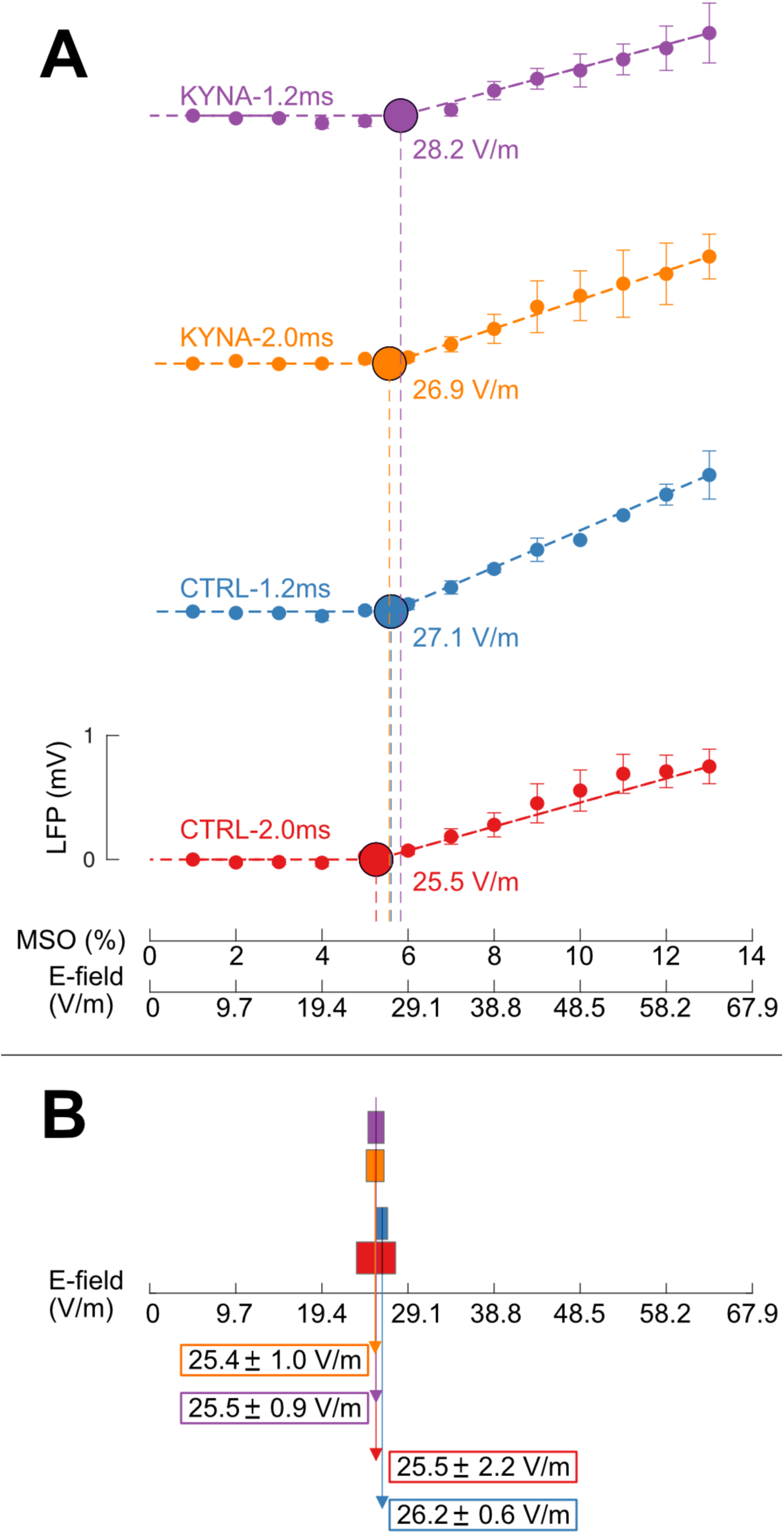
(A) E-field threshold curves in normal (CTRL) and KYNA saline in a single preparation. Colors indicate spike component and saline condition: Blue (CTRL-1.2 ms), Red (CTRL-2.0 ms), Purple (KYNA-1.2 ms), Orange (KYNA-2.0 ms). In each case, corresponding LFP amplitude at 500 µm depth is shown as a function of MSO (%). The MSO (%) was converted to E-field for the lowest threshold component using the threshold from Figure 7 (25.5 V/m). **(B) Thresholds for the 1.2 ms and 2.0 ms based on group-level (N=5) data in CTRL and KYNA.** In KYNA, the thresholds for the two spike components are statistically indistinguishable.

## DISCUSSION

### Toward a Mechanistic Understanding of TMS-Induced Neuronal Activation

Our findings are relevant to ongoing debates regarding the basic biophysical mechanisms of TMS. In our preparation, TMS could potentially activate four distinct neuronal populations: (1) the ascending segment of the climbing fibers, (2) Purkinje cells, (3) the ascending axons of the granule cells, and (4) Golgi cells.

As noted in the Introduction, an ongoing debate concerns whether TMS activates axons directly and if so, whether the activation is initiated at the terminals or at a bend. Based on the CSD profiles of the spike at 1.2 ms, we concluded that TMS activated the ascending segment of the climbing fibers with the current sink near the bend and the source in the axonal terminals. This conclusion is consistent with empirical studies of magnetic stimulation of isolated straight and bent nerves^40^, which showed reduced excitation thresholds at axonal bending sites, as well as a prior simulation-based hypothesis^76^. This, however, stands in contrast to some other views regarding how TMS may activate afferent axons in the cortex^19–22^.

Another outstanding issue is whether TMS can directly activate principal neurons and, if so, where the current is initiated. Our CSD analysis showed that TMS can directly activate Purkinje cells. For the spike at 1.2 ms, the current sink peaked at 420 μm in CTRL and 375 μm in KYNA; the slight shift of the peak toward the granular layer due to KYNA indicates that the climbing fibers partially contributed at this latency. In contrast, the current sink for the 2.0 ms spike component peaked at 485 μm (Purkinje cell layer) in both CTRL and KYNA. -- the indistinguishable CSD profiles confirming that climbing fiber activation was negligible at this latency. Since synaptic transmission is blocked in KYNA, the CSD profile for the 2.0 ms component in this condition must arise exclusively from currents in Purkinje cells directly activated by TMS. While the spatial resolution of our CSD analysis precludes a more precise localization, the current sink in the Purkinje cell layer most likely originates in the axon initial segment (AIS) – the established site for action potential initiation in Purkinje cells, owing to its high density of low-threshold voltage-gated Na^+^ channels^77–80^. When synaptic transmission is intact in CTRL, climbing fibers synaptically activate the proximal dendritic trunk of Purkinje cells as is well established. Accordingly, the CSD profile at 2.0 ms showed a current sink in the region of the proximal dendrites, but the peak of the sink was located in the region of the soma/AIS. Since the sink for the 2.0 ms component peaked at the same location in CTRL and KYNA, the CSD profile suggests that synaptic depolarization of the dendrites electrotonically spread to depolarize and activate the Purkinje cell AIS -- just as when synaptic transmission was blocked.

Although we obtained clear evidence for activation of the climbing fibers and Purkinje cells, there was no evidence that the remaining two types of neurons - the ascending granule cell axons and the Golgi cells - were activated even at the maximum E-fields produced by the clinical TMS stimulator used in our setup (∼250 V/m). Had TMS activated granule cells, the resulting parallel fiber activation would have produced the triphasic responses characteristic of parallel fiber activation in the molecular layer^70^. We hypothesize that this difficulty in activating the granule cells reflects thin diameters (0.88 µm and 0.64 µm^66^) of ascending granule cell axons and parallel fibers, which can substantially increase their activation thresholds^81,82^. The fast rise time and short duration of TMS pulses in the clinical TMS system used in this study, typical of clinical TMS systems^1,83^, may be responsible for this difficulty. This issue should be investigated further using TMS stimulators with controllable pulse parameters^84^, as it bears directly on the fundamental question of whether TMS can preferentially activate distinct neuronal populations.

In summary, the CSD analysis identified three types of currents that were activated by TMS: (1) CF_dir_ -- currents generated by direct, non-synaptic activation of the ascending climbing fiber segment, (2) PC_syn_ -- synaptic currents in Purkinje cells driven by TMS-activated climbing fibers, and (3) PC_dir_ -- currents generated by direct, non-synaptic activation of Purkinje cells by TMS. We hypothesized that the CSD profile of the spike at 2.0 ms was due to a mixture of PC_syn_ + PC_dir_ in CTRL and the profile in KYNA solely due to PC_dir_. A similar analysis for the spike at 1.2 ms suggested that this component was generated by a mixture of CF_dir_ + PC_syn_ + PC_dir_ in CTRL, and by a mixture of CF_dir_ + PC_dir_ in KYNA.

The slow wave with a peak at around 5 ms was produced by synaptic activation of the Purkinje cells by the climbing fibers since this component was completely abolished in KYNA. The CSD profiles revealed that the current sink underlying this wave peaked in the proximal trunk of the Purkinje cells, confirming that the current generator was located in the proximal dendritic trunk, with a corresponding source in the molecular layer. Figure S7 provides a schematic illustration of our hypothesis of the underlying currents for the spike and wave in CTRL and KYNA.

### E-Field Thresholds for TMS Activation of Specific Neuronal Populations

Another outstanding issue concerns the E-field thresholds required to activate afferent fibers, efferent fibers and cortical neurons differentially. This study provides, to our knowledge, the first measurements of thresholds for identified afferent fibers and cortical neurons. We were able to estimate the thresholds for specific neuronal populations by linking the empirically determined activation thresholds for the LFP spike to the underlying activation of climbing fibers and Purkinje cells identified with the CSD analysis. The E-field threshold for PC_dir_ corresponds to the threshold of the spike at 2.0 ms in KYNA (25.4 ± 1.0 V/m) since this component appears to be solely due to PC_dir_ in KYNA. The threshold for CF_dir_ can be estimated from the threshold of the 1.2 ms spike in KYNA, since the currents underlying this component reflect the mixture of CF_dir_ + PC_dir_. The threshold at 1.2 ms in KYNA was 25.5 ± 0.9 V/m. This threshold should have been lower than the 2.0 ms threshold in KYNA if the CF_dir_ threshold were lower than that for PC_dir_. Since they are virtually the same, the threshold for CF_dir_ must be equal to or higher than that for PC_dir_. Although it is not possible to directly estimate the threshold for CF_dir_, it appears to be similar to that for PC_dir_ because the slope of the LFP vs E-field curve for the spike at 1.2 ms in Figure 8 is significantly steeper in CTRL than in KYNA. According to our CSD analysis summarized above, the currents underlying the spike at 1.2 ms are PC_dir_ + CF_dir_ + PC_syn_ in CTRL and PC_dir_ + CF_dir_ in KYNA. Since PC_syn_ is generated by CF_dir_, if CF_dir_ is activated near the threshold of PC_dir_ and synaptically contributes to the LFP, the slope of the curve should be steeper in CTRL than in KYNA – as confirmed statistically. At 2.0 ms, the slope was steeper in CTRL compared to KYNA, but the difference was not statistically significant, perhaps due to relatively weak contribution of PC_syn_. Although indirect, our results suggest that the climbing fibers as afferent axons and Purkinje cells as principal cortical neurons can be activated at comparable thresholds.

Our estimated thresholds are consistent with those from previous studies using applied E-fields to directly stimulate neurons in the turtle cerebellum^58–60^. For a slow varying sinusoidal E-field (0.05 Hz-1.0 Hz) applied to the turtle cerebellum, the thresholds were reported as 10-20 V/m^58^. These values are lower than our estimates, but consistent with the fact that they are based on steady-state E-fields. The higher threshold in our experiments was likely due to the rapidly changing E-field (rise time 84 µs, pulse width 410 µs) since stimulus duration is a critical parameter in magnetic stimulation^85^.

The thresholds of ∼25 V/m for climbing fibers and Purkinje cells are slightly lower but close to those estimated in human studies - 33-50 V/m based on evoked potentials measured on the scalp with EEG^23,24^ and 50-70 V/m based on resting motor thresholds determined with EMG or with visual movement of a thumb^25–27^. The thresholds based on EMG are close to the TMS threshold for activating pyramidal neurons in the human M1^86^. However, it is still not known whether the 50-70 V/m thresholds measured with EMG reflect direct activation of corticospinal motor neurons or indirect activation via afferent axons synapsing onto them. TMS applied to M1 can activate multiple neuronal elements – axons, excitatory neurons, and inhibitory neurons – at the same applied E-field in non-human primates^38^. Thus, some neurons may be activated at E-fields lower than the resting motor threshold. This may explain the lower threshold found by some using evoked EEG response to estimate the thresholds^24^. Unlike these human studies, we measured the threshold for specific afferent fibers and cortical neurons. The difference between our lower estimates and those for humans may also be due to key procedural differences. The thresholds estimated in our study were obtained in an optimal condition with the E-field aligned to the longitudinal axis of stimulated axons or cell bodies. In human studies, such controls are difficult to achieve; thus, the stimulation was most likely suboptimal, leading to higher estimates. Moreover, the thresholds in human studies reflect the E-field values above the threshold criterion used in our study – that is, sufficient to evoke an LFP response significantly above the pre-stimulus noise level.

It is interesting to note that the estimates for the climbing fibers are similar to those obtained for peripheral nerves (25 V/m)^42,87^. Although this agreement is most likely coincidental, it suggests that the thresholds we estimated may reflect general properties of neuronal membranes with voltage-sensitive sodium channels. It is worthwhile to explore whether the activation thresholds across different methods and preparations can be related within a unified biophysical framework.

While we could relate our estimated thresholds to previous empirical estimates, the values are quite low compared to those obtained from simulation studies using realistic neurons (175-430 V/m)^20–22^. One contributing factor may be the difficulty of accurately modeling the extracellular microenvironment^31^, as most simulation studies are based on isolated neurons embedded in homogeneous saline environments^20–22,31^ – conditions that differ substantially from the *in vivo* case. Our study provides additional motivation for the on-going effort to resolve this wide discrepancy between the empirical human studies and the simulation studies^19,31,88^.

## CONCLUSIONS

In conclusion, our study provides new information on some of outstanding issues on the biophysical bases of TMS activation of neurons in the brain. We found that TMS can directly activate the ascending segment of the climbing fibers with the activation originating near the bend and intracellular currents directed toward the axonal terminals. We also found that TMS can directly activate the Purkinje cells, which are the principal neurons in the cerebellum, with the activation occurring in the soma region, probably arising from the initial segment, and intracellular currents directed toward the dendrites. We also showed that the climbing fibers and Purkinje cells can be activated directly by TMS at comparable thresholds. The thresholds for activating the climbing fibers and Purkinje cells were slightly higher than previous estimates for Purkinje cells in turtle based on direct application of a steady state electric field, but slightly lower than those previously estimated for neurons in the human primary motor cortex. However, there is a large discrepancy between our empirical estimates and those based on simulation studies using realistic neurons. We believe that the cellular mechanisms of TMS activation of the climbing fibers and Purkinje cells will help advance our understanding of biophysical bases of TMS, possibly widening applications of TMS-based neuroscience research.

## Abbreviations

CSD: current source density
CTRL: control normal physiological saline in bath
E-field: Induced electric field
EEG: Electroencephalography
EMG: Electromyography
KYNA: kynurenic acid
LFP: local field potential
M1: primary motor cortex
MEP: motor evoked potential
TMS: transcranial magnetic stimulation

## Acknowledgments

This work was supported by the NIH Brain Initiative 1R01NS112183, the NIH Center for Mesoscale Mapping P41EB030006, and NIH R01MH130490. The authors would like to thank the Martinos Center TMS Core for help in the initial stages of this work, and Dr. Giorgio Bonmassar (Martinos Center for Biomedical Imaging, Department of Radiology, Massachusetts General Hospital, Boston, MA) for help with construction of TMS coils during the course of this work.

