## Supplementary Figures and Methods for "Cellular Mechanisms of Transcranial Magnetic Stimulation in Climbing Fibers and Purkinje Neurons in the Cerebellum"

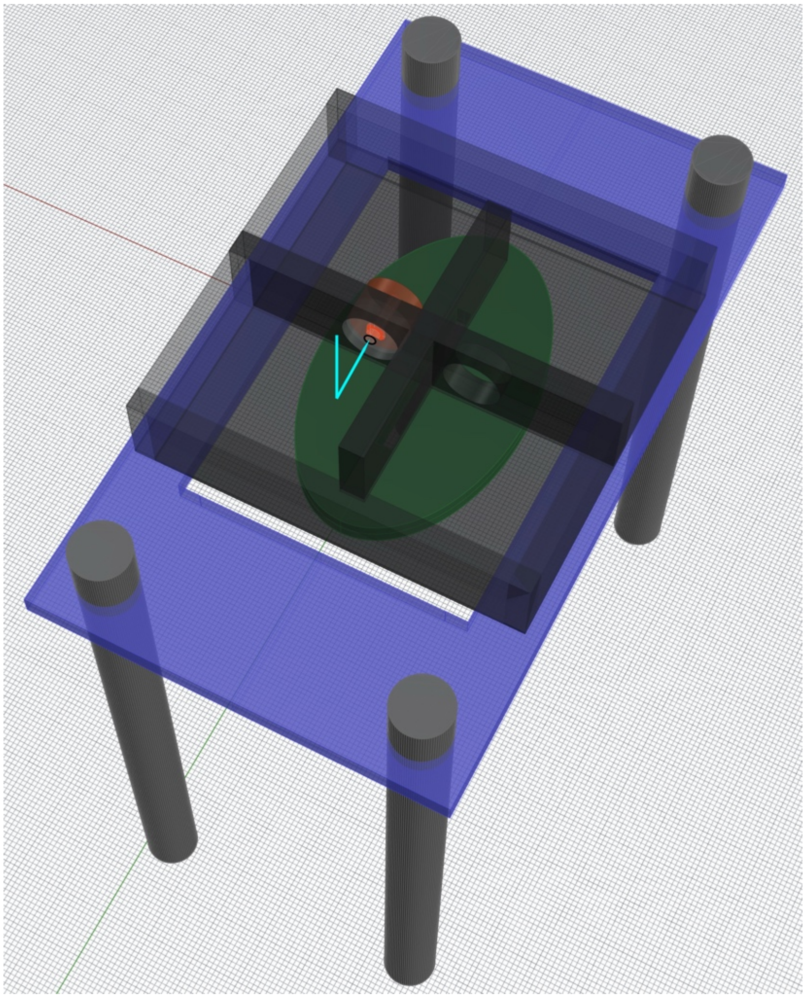

**Fig. S1.** Setup showing the current concentrator recording chamber placed on a 3d-printed plastic support (purple) placed on 4 brass rods screwed into the table. The coil surface was 3 mm below the chamber and mounted on a separate 3d-printed support screwed into the table. The glass pipette was bent at 90 degrees and advanced into the cerebellum which was placed inside the tunnel.

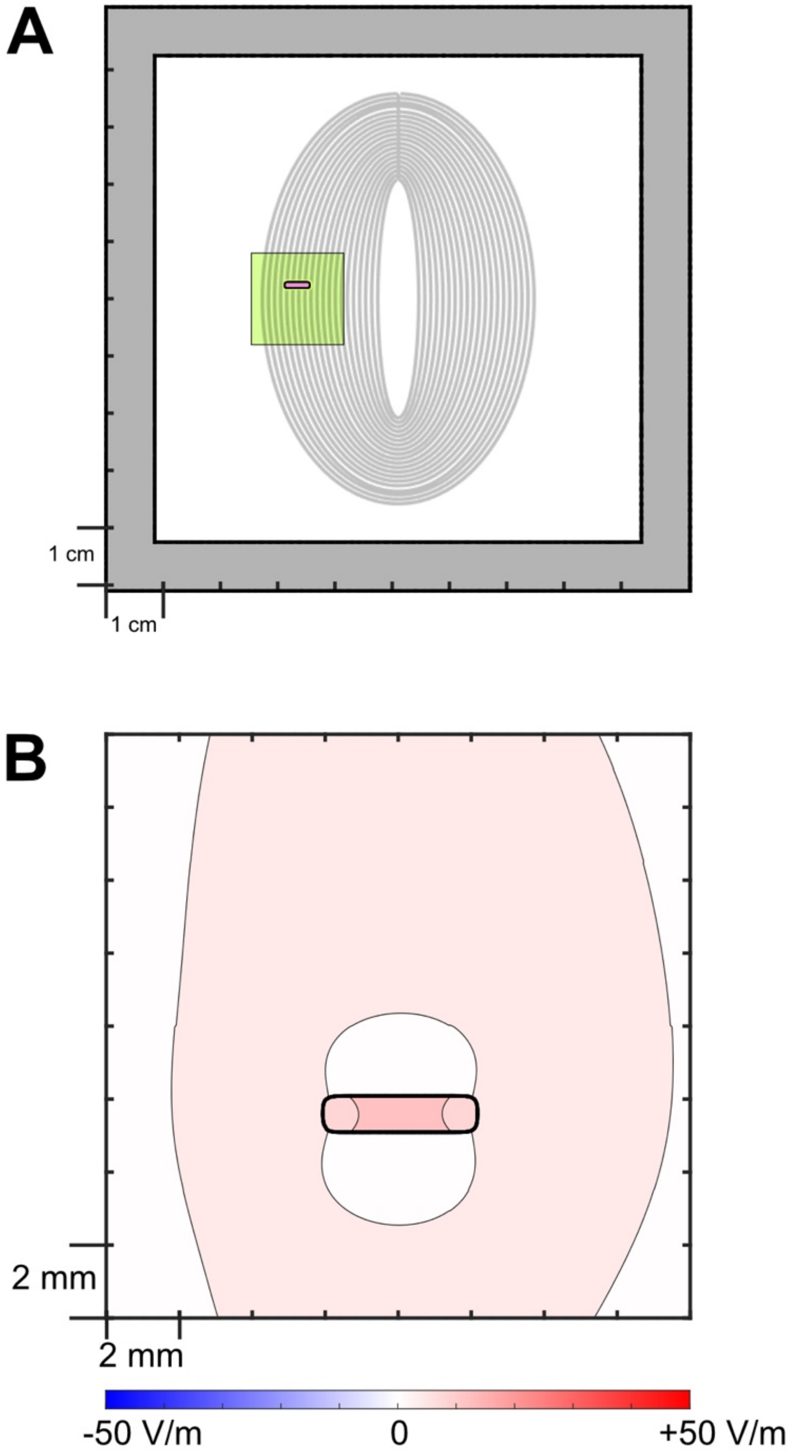


**Fig. S2.** In the absence of the current concentrator (Figure 1A-D), the E-field in the tissue is a factor of 3.2 smaller. Based on BEM-FMM simulations of the realistic setup geometry, we expect only 13 V/m in the tissue at MSO 5%. In comparison, with the current concentrator design, we can expect 41.7 V/m at MSO 5% (see Figure 1D).

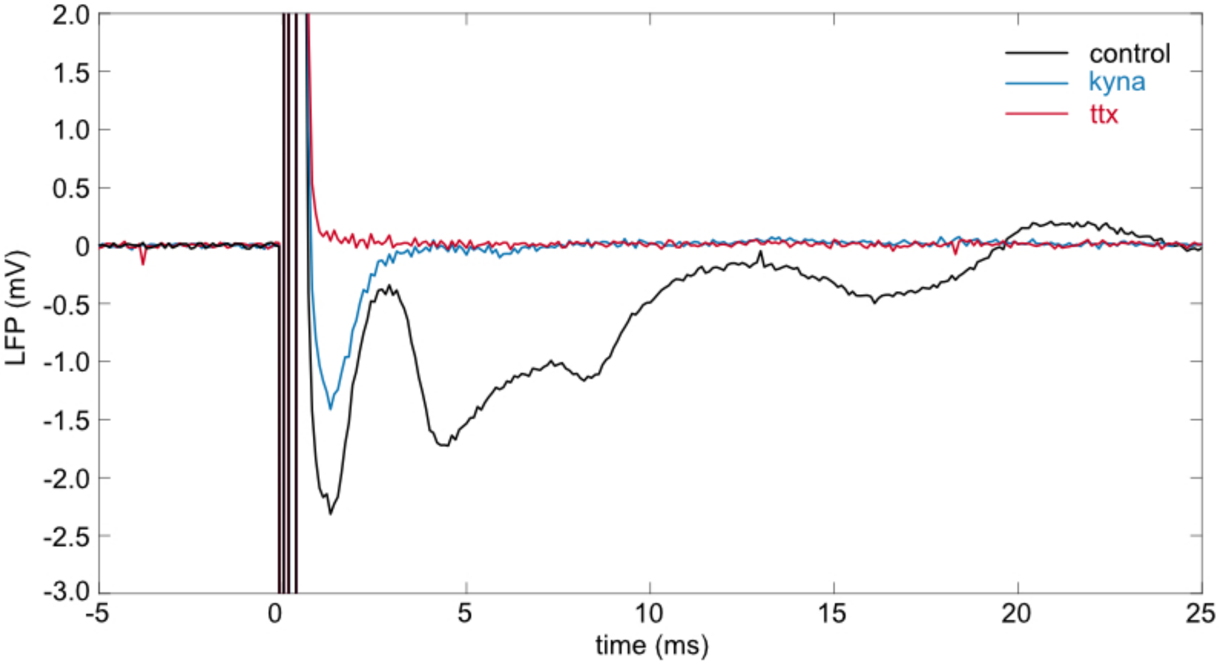


**Fig. S3.** Additional example to Figure 1H. Extracellular LFP evoked response to TMS measured 500 μm from the ventral surface in the PC layer in the control condition with normal physiological saline (black), when excitatory synaptic transmission was blocked with 10 mM KYNA (blue), and when voltage-gated $Na^{+}$ channels were blocked using 2 μM TTX (red). TMS electric field in the tissue was suprathreshold. Each plot is average of 3 trials.

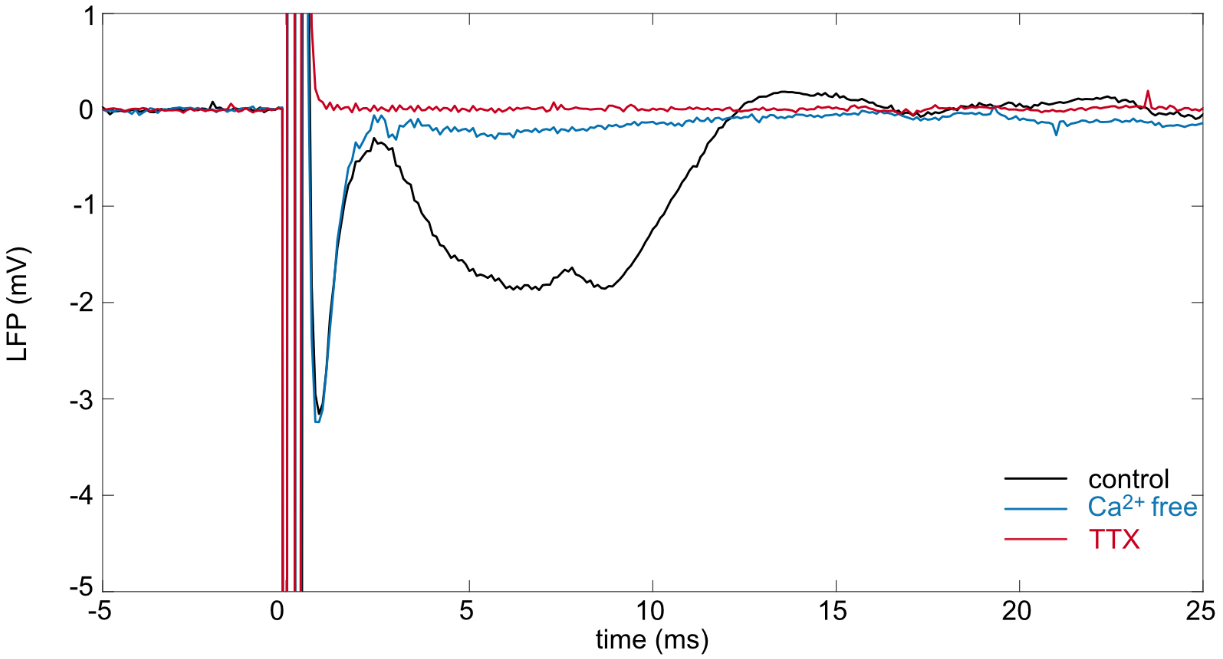


**Fig. S4.** Extracellular LFP evoked response to TMS measured 500 μm from the ventral surface in the PC layer in the control condition with normal physiological saline (black), in $Ca^{2+}$ free Ringer with $Mn^{2+}$ replacement (blue), and when voltage-gated $Na^{+}$ channels were blocked using 2 μM TTX (red). TMS electric field in the tissue was suprathreshold. Each plot is average of 3 trials.

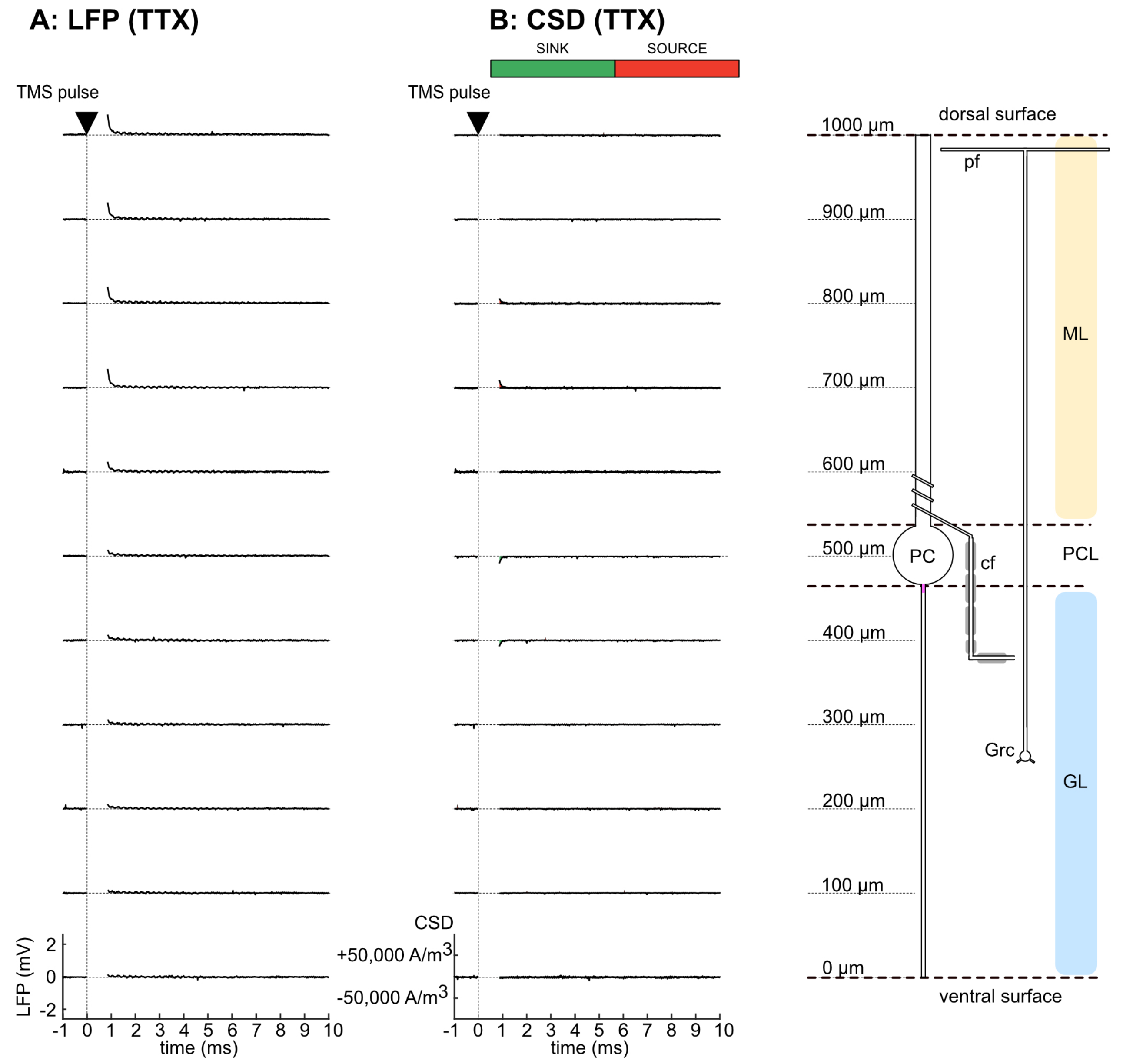


**Figure S5.** Laminar LFPs and CSD due to a single TMS pulse in Na+ channels blocked by application of TTX (2 μM). (Same preparation as Figures 2, 3).

(A) Laminar profile of evoked LFPs (10 ms window post-TMS).

(B) CSD profile of the responses in (A) assuming tissue conductivity of 0.2 S/m.

Cell layers are indicated on the right (ML: molecular layer, PCL: Purkinje cell layer, GL: Granular layer). PC: Purkinje cell, Grc: Granule cell, cf: climbing fiber, pf: parallel fiber. Data shown are for same preparation as Figures. 2, 3.

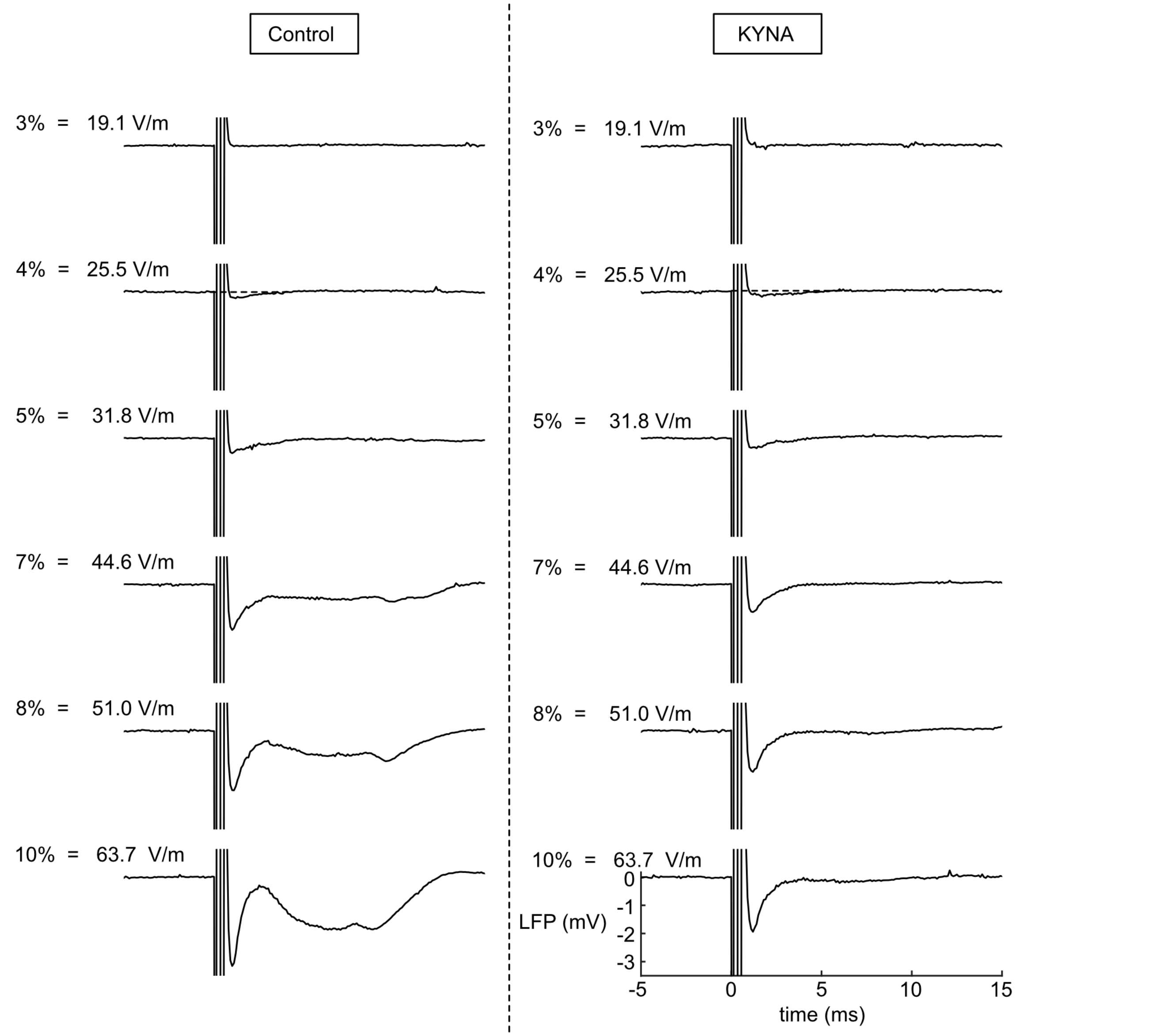


**Figure S6.** LFP in the PC layer (500 μm from ventral surface) as a function of the TMS intensity in normal saline vs when synaptic transmission was blocked (10 mM KYNA).

The left column shows the effect of TMS intensity on the LFP in the PC layer in normal saline and shows similar behavior to the preparation in Figure 6. The right column the effect of TMS intensity on the LFP at the same location when synaptic transmission was blocked. The slow 5 ms component is abolished. The initial spike is affected by KYNA application however the threshold of the initial component is very close that in normal physiological saline. Since this experiment did not have a calibration, to convert MSO to E-field, we used the E-field threshold from the calibrated experiments (25.5 V/m).

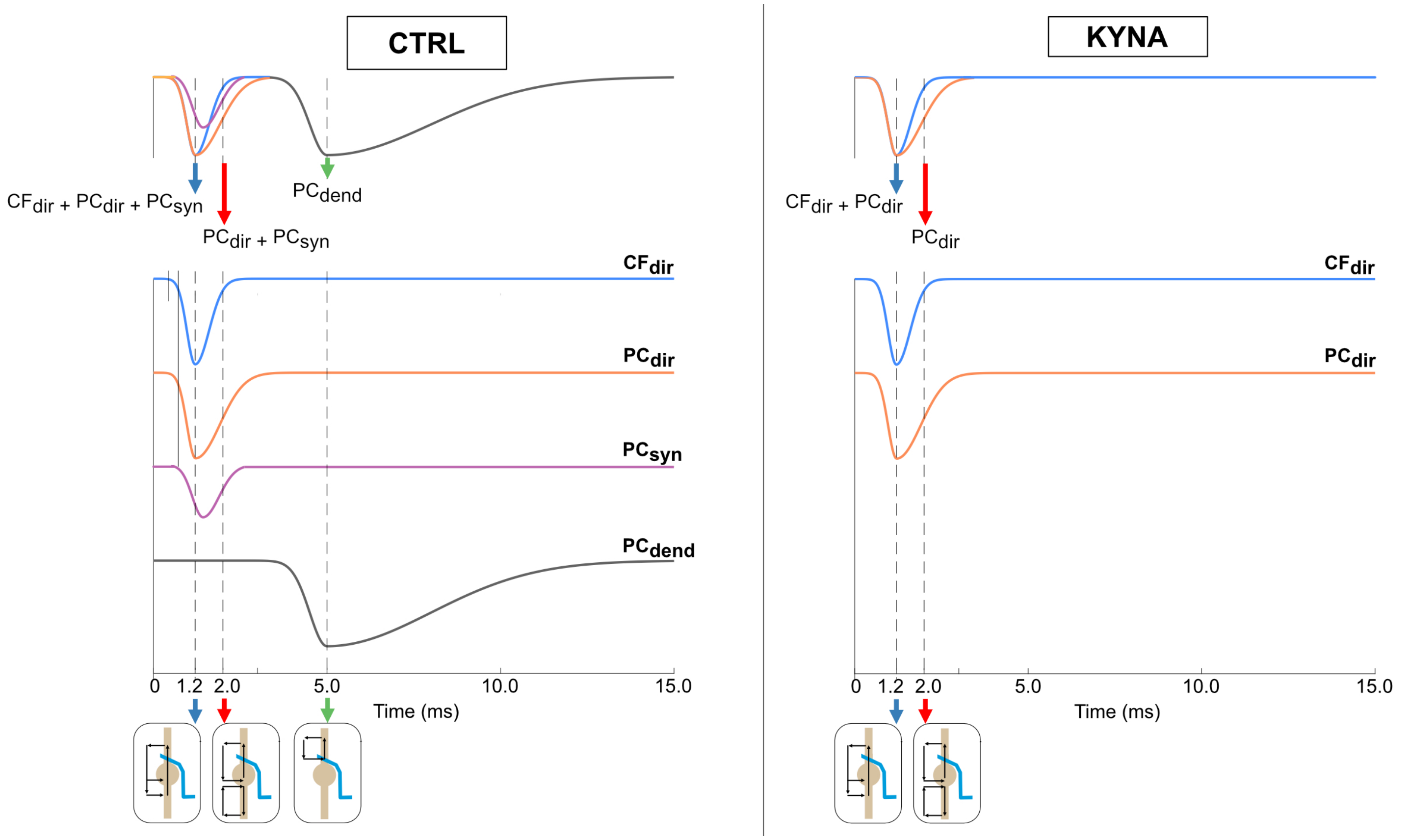


**Figure S7.** Simplified schematic representation to explain the potential generators of the currents underlying the 1.2 ms and 2.0 ms components in normal (left) and KYNA (right) saline.

$CF_{dir}$ = non-synaptic direct activation of climbing fiber

$PC_{syn}$ = synaptic activation of Purkinje cell by the climbing fiber

$PC_{dir}$ = non-synaptic direct activation of the Purkinje cell.

$PC_{dend}$ = synaptically activated dendritic currents in the Purkinje cell

In (A), the 1.2 ms component is a mixture of $CF_{dir}$, $PC_{syn}$ and $PC_{dir}$. The 2.0 ms component is a mixture of $PC_{syn}$ and $PC_{dir}$.

In (B) with synaptic transmission blocked by KYNA, the 1.2 ms component is a mixture of $CF_{dir}$ and $PC_{dir}$. The 2.0 ms component is $PC_{dir}$.

### Methods

#### Turtle Cerebellum

We used an isolated, intact, whole turtle cerebellum submerged in physiological saline (Ringer's solution). The cerebellum of turtle is attached to the brainstem with two cerebellar peduncles located behind the optic tectum. The cerebellum is a lissencephalic structure *in vivo*, but it can be flattened into a disk *in vitro*. Though the turtle cerebellum is not foliated like in humans, its basic local cellular organization is evolutionarily conserved[1] . The turtle cerebellum is a 3-layer structure with a thickness of 1 mm and 4-5 mm wide along the mediolateral and the rostrocaudal directions. It consists of the molecular layer in the dorsal half and the granular layer in the ventral half. The granular layer (the ventral 450 µm of the cerebellum contains the granule cells, which send their unmyelinated axons (0.88 µm diameter) up into the molecular layer where they divide like the letter T into two 1.5 mm-long branches that run along the mediolateral axis. The parallel fibers (~0.64 µm diameter) make up the bulk of the molecular layer (dorsal 450 µm of the cerebellum) and are largely unmyelinated though they become myelinated deeper into the molecular layer close to the Purkinje cell layer. At ~500 µm depth, the Purkinje soma (~ 30 µm diameter) form a 1-2 cell thick layer on top of the granular layer. The Purkinje cells extend elaborate dendritic arbors into the molecular layer; the dendritic arbor is flattened in the parasagittal plane at right angles to the parallel fibers making the major neural elements cerebellar cortex mutually orthogonal to each other. The climbing fiber bends when it enters the granular layer and is a thick (1.13 µm diameter) myelinated axon. About the level of the Purkinje cell soma, the climbing fiber loses its myelin sheath and starts to climb the dendritic arbor of the Purkinje cell. In the case of the turtle cerebellum, the climbing fiber stays in the ventral 1/3 of the molecular layer. Each Purkinje cell is furnished with a single myelinated climbing fiber.

This preparation is ideal for study of the biophysical mechanisms underlying TMS for several reasons: (1) The entire cerebellum *in vitro* can be kept physiologically functional for extended periods since it can withstand hypoxia, (2) unlike slice preparations, the normal cellular circuit is completely intact, and (3) the flat disk-like geometry combined with the mutually orthogonal arrangement of neural structures in the cerebellum allow the TMS electric field to be applied along specific axes (Fig. 1C-D).

#### Tissue Preparation

The surgical procedure involved in the preparation of the turtle cerebellum was approved by the Institutional Animal Care and Use Committee (IACUC) at our institution. The brain of the red-eared slider turtle (*Pseudemys Scripta Elegans*) was removed rapidly after decapitation and craniotomy and placed in cold artificial cerebrospinal fluid (Ringer's solution) (100 mM NaCl, 5 mM KCl, 40 mM $NaHCO_3$, 2.5 mM $CaCl_2$, 1.25 mM $MgCl_2$, 20 mM D-glucose). The entire cerebellum including the cerebellar peduncles was dissected away from the brainstem and the optic tectum under a microscope. The peduncles were further trimmed away leaving behind just the intact isolated cerebellum.

**TMS Coil and Stimulator**

We constructed a circular TMS coil by winding high-temperature magnet wire (18 AWG, Remington, 19 turns, 1 layer, 7.2 cm x 5.8 cm size). The coil was coated with epoxy (834B, MG Chemicals, Burlington, Ontario, Canada). The coil inductance was 18.3 $\mu$H and its resistance was 300 m$\Omega$. The coil was connected to a commercial TMS stimulator (MagPro R30, MagVenture, Farum, Denmark). Throughout all studies, we only used single-pulse TMS stimulation. The pulse width was 410 $\mu$s and the rise time was 84 $\mu$s (Fig. 1E). In all the experiments, the electric field was oriented from the dorsal to the ventral surface (i.e., along the length of the Purkinje neurons, pointed from the dendrites towards the soma) (Figs. 1C-D).

**Recording chamber and Tissue Placement**

We used a current concentrator design (Figs. 1A-B**)** inspired by a similar design used for direct current stimulation[2] . The acrylic chamber was 9.5 cm x 9.5 cm in size (Fig. 1A). The intact cerebellum was placed inside the 1.2-cm-diameter tunnel which had a 3-mm-diameter opening. The cerebellum was surrounded by vacuum grease (DuPont Molykote High-Vacuum Grease, DuPont Inc., USA) and then kept in place with a plastic insert which also had a 3-mm-diameter hole. Fig. 1B shows the setup. The chamber was then filled with Ringer's solution. The chamber was placed on top of a 3d-printed platform (Fig. S1). The platform was held up using four brass rods that were screwed into the top of the anti-vibration table. The coil was also bolted using set screws to a 3-d printed support, which was also screwed to the top of the table. The coil and its support were directly below the chamber but did not make any physical contact with the chamber. We measured a 2.5 mm gap between the surface of the TMS coil and the bottom of the recording chamber.

**Recording Local Field Potentials in the cerebellum during TMS**

The extracellular LFPs in response to TMS were recorded in the tissue using a glass micropipette (5-10 µm tip diameter, GC150F-10, Harvard Apparatus), filled with 150 mM NaCl (1-5 M$\Omega$ impedance). The pipette positioning was handled with a precision motorized micromanipulator (MP-285, Sutter Instruments, CA). The micropipette was connected to the headstage of an extracellular recording amplifier (A-M Systems Inc.,

Model 1800 Microelectrode AC Amplifier, Sequim, WA, USA). The measurements were differential measurements. Both reference and ground electrodes were fixed in the bath. The signal wire was inserted into the saline in the glass micropipette just deep enough to make contact. The signal was amplified (100x) and filtered (0.1 Hz, 5000 Hz). The amplifier output was digitized using in-house software using a DAQ device (National Instruments USB-6263). The sampling rate was 50 kHz. The recordings were then analyzed using in-house software written in MATLAB (Mathworks, Natick, MA).

**Pharmacological studies**

For N=5 animals, we recorded the LFPs at (v-500) μm in the Purkinje cell layer in 3 conditions: (1) Control, (2) KYNA (excitatory amino acid-mediated synaptic transmission blocked 30 min after application of 10 mM kynurenic acid), (3) TTX (block of voltage-gated Na+ channels with 2 μM tetrodotoxin). For a fixed stimulus intensity, we recorded 3 trials in each condition without moving the recording electrode.

**Laminar Profile and Current Source Density Analysis**

For N=5 animals, we recorded the laminar profile of the TMS-evoked LFPs by positioning the glass micropipette at different depths in the cerebellum (every 100 μm starting 300 μm from the ventral surface in the saline, passing through the tissue, to 300 μm from the dorsal surface in the saline). At each depth, 2 trials were recorded. In all cases, the glass micropipette was coated with Sylgard (Sylgard 184, Dow) to reduce the electrode capacitance. Once the control condition laminar profiles were recorded, we added 10 mM KYNA to block excitatory amino acid-mediated synaptic transmission and recorded the laminar profile again in the KYNA condition. At the end of each of the experiments, we added 2 μM TTX (block of voltage-gated Na+ channels) to the bath to verify that there was only the TMS pulse artifact remaining.

The laminar profiles for both control and KYNA conditions were plotted as a function of depth for study. We used in-house MATLAB-based scripts for analysis. We calculated the current source densities (CSDs in Amperes/$m^3$) from the second spatial derivative of the recorded LFPs using the approach in [3,4]. At any depth r, the CSD at the depth is given as,

$$CSD(r) = -\sigma D_4(r)$$

where $\sigma$ is the extracellular conductivity (0.2 S/m) and $D_4(r)$ is the smoothed second derivative at r. $D_4(r)$ was computed using the 4-point formula in [3,4].

$$D_4(r) = \frac{1}{kh^2} \sum_{m=-3}^{+3} (a_m \phi(r + mh))$$

where $a_0$ = -20, $a_{\pm1}$ = -5, $a_{\pm2}$ = 6, $a_{\pm3}$ = 9, and h = 100 µm.

After studying the features of the TMS-evoked response, the CSD profile corresponding to components of interest were plotted in both the control and the KYNA condition for comparison.

**Determination of TMS Thresholds for Different Components using Electric Field Calibration**

For N=5 animals, we determined the electric field thresholds for different components in the TMS response (1.2 ms, 2.0 ms and the slow wave at 5 ms) at (v-500) µm in the Purkinje cell layer. To do this, we first calibrated the recording chamber in each of the N=5 cases. Empirical calibration of the chamber accounts for experiment-to-experiment variability in the induced electric field in the tissue for a given stimulus intensity and yields a conversion factor of MSO (in %) to electric field in the tissue (V/m). The TMS-induced electric field in the tissue was experimentally determined by recording the electric potential along a line starting in the saline, entering the cerebellum at the ventral surface and ending at the dorsal cerebellar surface in the tunnel. The potential was recorded every 250 or 500 µm in the saline and every 50 or 100 µm in the tissue (N=5 animals) at MSO 5% (1.5 A/µs). The glass micropipette (5-10 MΩ impedance, GC150F-10, Harvard Apparatus) was pulled and then coated with Sylgard (Sylgard 184, Dow) to reduce the electrode capacitance. The pipette was connected to the headstage of an intracellular amplifier (IE-210, Warner Instruments, Hamden, CT). The signal was amplified (10x) and not filtered. The data were digitized using a DAQ device (National Instruments USB-6263) and in-house recording software. The sampling rate was 500 kHz. At each location, we acquired 10-20 trials. These recordings were done inside an electromagnetically shielded room with the setup placed on an acrylic table to prevent corruption of the recorded potentials. The data were analyzed using in-house MATLAB software. At each location, the average peak electric potential was determined. For our coil and TMS stimulator, the peak was reached ~6 µs after the stimulator was triggered (Fig. 1E). The line profile of the average peak electric potential was plotted in saline and tissue as a function of distance. To determine the electric field from the electric potential, we performed a least-squares line fit using MATLAB (Mathworks, Natick, MA). This gave us the empirically determined electric field in the saline and inside the cerebellum at stimulator intensity of 5%. For each case (N=5), we noted the calibration factor (V/m per %) specific to that experiment.

Once the calibration was performed, we positioned the glass micropipette electrode (1-5 MΩ impedance) at (v-500) µm in the Purkinje cell layer and slowly varied the stimulus intensity from low to high and recorded 3 extracellular LFPs at each intensity with the electrode position fixed. Typically, the stimulus intensities used were 2, 3, 4, 5, 6, 8, 10, 15, 20%. The LFPs were recorded with the same intracellular amplifier (IE-210, Warner Instruments, Hamden, CT). The signal was amplified (10x) and filtered (0.1 Hz, 5000 Hz).The data were digitized using a DAQ device (National Instruments USB-6263) and in-house recording software. The sampling rate was 50 kHz. For each of the N=5 studies, we determined the average LFP amplitude for each of the response components (1.2 ms, 2.0 ms, 5.0 ms) as a function of stimulus intensity (in %). Using the calibration factor specific to each experiment, we were able to obtain the response amplitudes as a function of the experimentally measured TMS-induced electric field in the tissue. The LFPs corresponding to each component were normalized and then averaged across N=5 and then plotted as a function of the electric field in the tissue. Using line fitting, the electric field threshold corresponding to each component was determined.

For a different set of N=5 animals, we repeated the LFP vs stimulus intensity measurements in the control condition and then added 2 mM KYNA to the bath without moving the micropipette. Now with excitatory synaptic transmission blocked by the KYNA, we repeated the LFP vs stimulus intensity measurements in this KYNA condition. The recording parameters were same as for the previous studies. We used the threshold obtained from the studies with the calibrated chambers to convert the MSO to electric field in the tissue. We then plotted the peak LFPs corresponding to the 1.2 ms and 2.0 ms responses in the control and the KYNA condition and compared their electric field thresholds and their behavior.

#### Boundary Element Model (BEM) Simulation for Comparison to Empirical Measurement

We used the software toolkit *boundary element model – fast multipole method* (BEM-FMM)[5,6] to perform a simulation of the TMS-induced electric field in the cerebellum. Our TMS coil was modeled in the BEM-FMM software using 43K elementary current segments. We used dI/dt = 1.5 A/µs, corresponding to 5% maximum stimulator output (MSO) for the MagVenture MagPro R30 (MagVenture, Denmark) stimulator with our custom circular coil. The dI/dt value was recorded from the stimulator with 5% MSO. To perform the simulation, we used a geometric model of the chamber with the horizontal and vertical barriers and the tunnel containing the cerebellum held in place with the plastic insert and vacuum grease. The conductivity of the saline was assumed to be 1.33 S/m and that of the cerebellum was assumed to be 0.2 S/m (based on experimental result in

[7]). The region next to the cerebellum is a mix of vacuum grease and Ringer solution. We assumed this region to be 0.6 S/m accounting for some leak in the tunnel. The BEM-FMM algorithm is implemented in MATLAB (for both Windows and Linux) and available via a GitHub repository. All the meshes used in our simulation were 3d-printable mesh models constructed in Rhino (Robert McNeel & Associates, Seattle, WA). The triangle meshes were stored in the *stl* format and converted to MATLAB *mat* format for use in the simulations. As in the experiment, the coil model was placed 2.5 mm below the base of the chamber. We computed the total induced electric field in the tissue and the saline. We also computed the electric potential along a line along the y axis, starting in the saline and passing through the tissue. We plotted the line profile of the electric potential from the BEM simulation over that obtained from the experiments. We also compared the electric field predicted by BEM to that measured by us experimentally.

turtle cerebellum and the interpretation of current source-density analysis. Journal of Neurophysiology *72*, 742–753.
